# Specificity, length, and luck: How genes are prioritized by rare and common variant association studies

**DOI:** 10.1101/2024.12.12.628073

**Authors:** Jeffrey P. Spence, Hakhamanesh Mostafavi, Mineto Ota, Nikhil Milind, Tamara Gjorgjieva, Courtney J. Smith, Yuval B. Simons, Guy Sella, Jonathan K. Pritchard

## Abstract

Standard genome-wide association studies (GWAS) and rare variant burden tests are essential tools for identifying trait-relevant genes. Although these methods are conceptually similar, we show by analyzing association studies of 209 quantitative traits in the UK Biobank that they systematically prioritize different genes. This raises the question of how genes should ideally be prioritized. We propose two prioritization criteria: 1) trait importance — how much a gene quantitatively affects a trait; and 2) trait specificity — a gene’s importance for the trait under study relative to its importance across all traits. We find that GWAS prioritize genes near trait-specific *variants*, while burden tests prioritize trait-specific *genes*. Because non-coding variants can be context specific, GWAS can prioritize highly pleiotropic genes, while burden tests generally cannot. Both study designs are also affected by distinct trait-irrelevant factors, complicating their interpretation. Our results illustrate that burden tests and GWAS reveal different aspects of trait biology and suggest ways to improve their interpretation and usage.

## Introduction

A central goal of human genetics is to identify which genes affect traits and disease risk and to what extent. This is essential for addressing fundamental questions such as: What biological processes underlie trait variation? Which genes and pathways are most critical for understanding those processes? Which genes could serve as potential therapeutic targets?

While many techniques exist to study gene function in model systems or *in vitro* (e.g., [1–3]), the study of organism-level traits in humans largely relies on naturally occurring genetic variation, primarily through genome-wide association studies (GWAS) [4].

GWAS have been deeply informative about the genetic basis of complex traits, from uncovering actionable drug targets [5] to identifying trait-relevant cell types and programs [6–9]. However, it remains unclear how best to extract biological insight from GWAS results. First, GWAS do not directly pinpoint relevant genes, as most associated variants are non-coding [10]. Moreover, a surprisingly large fraction of the genome contributes to the heritability of many traits [11–13], and associated variants often cannot be mapped to genes with clear phenotypic relevance.

Recently, large whole-exome and whole-genome sequencing datasets have enabled the direct study of genes through rare protein-coding variants, which have typically been excluded or underpowered in GWAS [14]. To boost statistical power, these variants are analyzed using burden tests [15, 16]. Burden tests aggregate variants — typically loss-of-function (LoF) variants — within a gene to create a “burden genotype”, which is then tested gene-by-gene for association with phenotypes. This is similar to common-variant GWAS but focused on rare variants collapsed at the gene level.

Despite this conceptual similarity, recent work has found anecdotally that LoF burden tests and GWAS discover distinct genes, though with some overlap [17, 18]. In a systematic analysis, Weiner *et al*. found that burden tests appear less polygenic and tend to prioritize genes that are seemingly more closely related to trait biology [19].

To better understand these differences, we analyzed the results of GWAS and LoF burden tests for 209 quantitative traits in the UK Biobank [14, 16, 20]. We show that burden tests and GWAS prioritize different genes, and these differences cannot be explained by differences in power or challenges in linking GWAS variants to genes.

The discrepancy between GWAS and LoF burden tests raises thorny questions. By what criteria does each prioritize genes, and how do these relate to the underlying biology? Which method is more relevant for understanding trait biology? Which is better suited for downstream applications, such as drug target discovery?

We analyze association study results and use population genetics models to address these questions. Our results show that burden tests tend to prioritize trait-specific genes — those primarily affecting the studied trait with little effect on other traits — while GWAS also capture more pleiotropic genes often missed by burden tests. Additionally, we highlight the impact of trait-irrelevant factors on discovery, such as gene length and random genetic drift. Ultimately, GWAS and LoF burden tests reveal distinct but complementary aspects of trait biology, with important implications for interpreting and using association studies.

## Results

### LoF burden tests and GWAS prioritize different genes

GWAS and LoF burden tests are conceptually similar (Figure 1**A,B**), but previous studies have highlighted key differences in their findings [19]. To more thoroughly quantify how concordant these methods are in prioritizing genes and genomic loci based on p-values, we systematically compared GWAS and LoF burden test results for 209 quantitative traits from the UK Biobank [16] (Methods).

**Figure 1:**
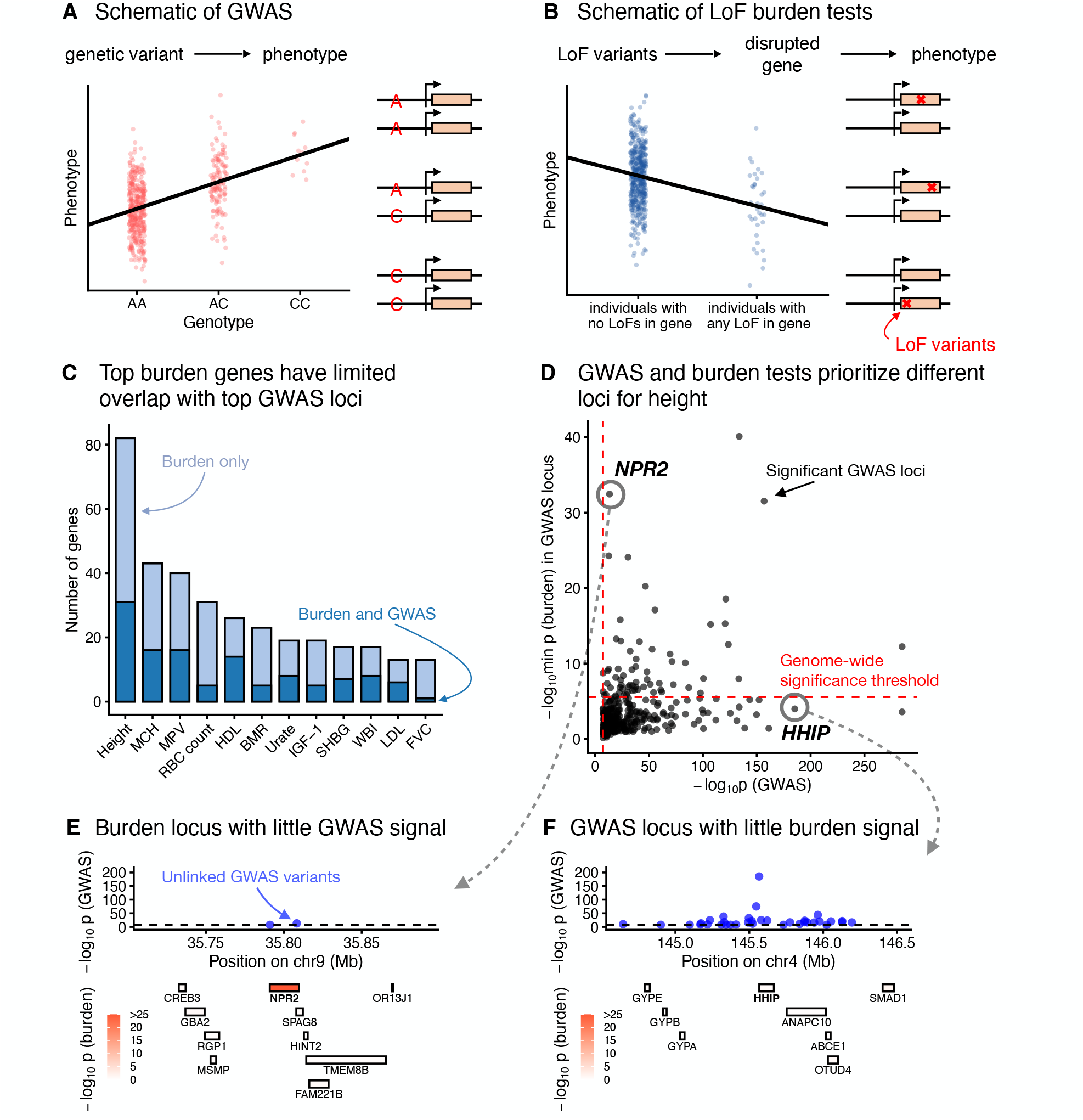
GWAS and LoF burden tests prioritize different loci. Schematics of **A**) GWAS and **B**) LoF burden tests. **C**) Bar charts of genome-wide significant LoF burden test genes, split by whether or not the gene overlaps a top GWAS locus. **D**) Minimum LoF burden test p-values for any genes overlapping a genome-wide significant GWAS locus plotted against the minimum GWAS p-value within the locus. **E**) The genomic region surrounding NPR2. Top panel: GWAS p-values of approximately independent genome-wide significant GWAS hits. Bottom panel: location of genes colored by LoF burden test p-values. **F**) Similar to **E** for the genomic region surrounding HHIP.

In principle, discordance between GWAS and LoF burden test results could be driven by technical artifacts. The causal genes driving GWAS hits are usually unknown, and errors in linking hits to genes could reduce the overlap between genes prioritized by the two study designs. Additionally, LoF burden tests typically discover fewer genes than GWAS. Hence, some discordance could potentially be driven by differences in power.

To minimize these technical effects, we maximized overlap whenever possible and controlled for power. We defined GWAS loci by taking a 1Mb window around each genome-wide significant GWAS hit and merging overlapping windows. We ordered these loci by the minimum GWAS p-value within the locus, and considered sufficiently significant loci “top GWAS loci”, with the significance threshold chosen so that there were an equal number of top GWAS loci and genomewide significant LoF burden test genes. Across traits we found that only 26% of significant LoF burden genes (480 out of 1852) are contained in a top GWAS locus (Figure 1**C**, Supplementary Figure S1, Methods).

Figure 1**D** shows the minimum LoF burden test and GWAS p-values for the 382 genome-wide significant GWAS loci for height. The results of the two study designs are somewhat concordant (Spearman’s *ρ* = 0.46), suggesting that they are not uncovering totally disparate axes of biology. Yet, there is little overlap in the top hits, with many significant GWAS loci not containing a single significant burden gene. This pattern is not unique to height (Supplementary Figures S2 and S3), and these results are robust to how we partition the genome into loci or if we look at signals in LD blocks instead of GWAS loci (Supplementary Figures S4–S7).

To further explore this lack of overlap, we considered two examples of discordantly ranked loci. Figure 1**E** shows the *NPR2* locus. *NPR2* is the second most significant gene in the LoF burden tests, but it is contained in the 243^rd^ most significant GWAS locus. That this locus is significant in both association tests is not surprising: mutations in *NPR2* have been linked to short stature in humans and mice [21–26]. Yet, hundreds of loci are more strongly prioritized by GWAS, including the *HHIP* locus (Figure 1**F**). The *HHIP* locus has numerous uncorrelated GWAS hits (*r*^2^ < 0.1) with p-values as small as 10^−185^. *HHIP* is a biologically sensible hit for height [27] as HHIP has been implicated in osteogenesis [28], and interacts with three different hedgehog proteins [29, 30], which are involved in body patterning and limb formation [31]. Nonetheless, there is essentially no burden signal for *HHIP* or any of the other genes in the locus. These differences motivated us to explore why GWAS and LoF burden tests might rank loci so differently.

### How should genes be prioritized?

Given the extensive differences in how GWAS and LoF burden tests rank genes, we are faced with an underexplored question: if we could precisely measure any possible quantity of interest for each gene, what properties would make us want to rank one gene higher than another for a given trait? That is, how should genes ideally be prioritized?

We propose two distinct properties by which one may wish to prioritize genes: *trait importance* and *trait specificity*. Imagine a gene that is only expressed in developing bones and whose disruption results in shorter stature but has minimal effects on other traits (Figure 2**A**). In some sense this is a quintessential “height gene”, and we might want this gene to be highly ranked in association studies. On the other hand, consider a broadly expressed transcription factor whose disruption results in an even greater reduction of height, but also disrupts the normal functioning of numerous organ systems. This is less obviously a “height gene”, but it has a larger impact on height than the first gene. We define trait specificity and trait importance such that the first gene has higher trait specificity, but the second gene has higher trait importance.

**Figure 2:**
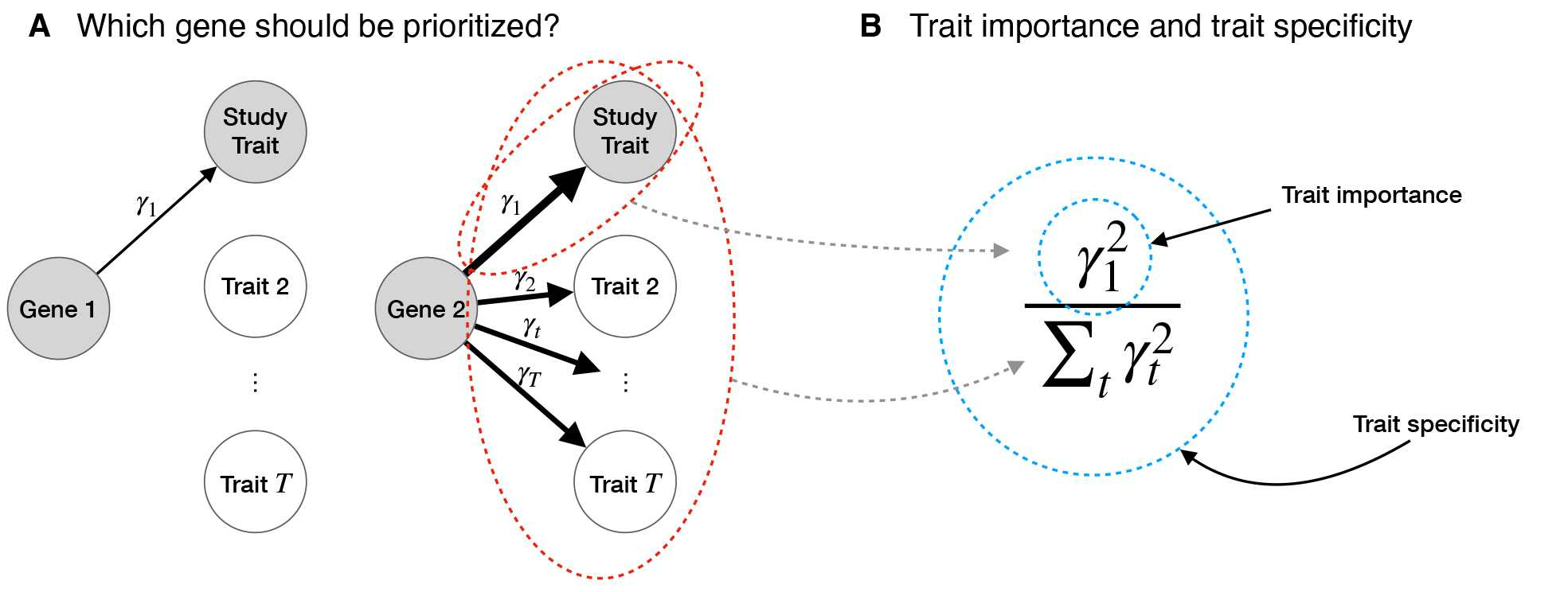
How should genes be prioritized? **A**) A cartoon of two genes that affect a trait under study. Widths of arrows represent relative effect sizes. Gene 1 is more trait specific but Gene 2 is more trait important. **B**) Formal definitions of trait importance and trait specificity for genes in the context of LoF burden tests. The effect of an LoF in the gene on trait t is γ_t_, with trait 1 being the study trait. We define trait importance as 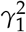 and trait specificity as 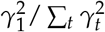.

Formally, we define the trait importance of a variant as its squared effect on the trait of interest, considering high-impact variants important regardless of the direction of their effect. We define the trait importance of a *gene* as the trait importance of LoFs in that gene. Throughout, we use *α*_*t*_ to refer to the effect size of a variant on trait *t*, and *γ*_*t*_ to refer to the LoF burden effect size of a gene, so trait importance for trait 1 would be 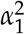 and 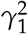 for variants and genes respectively. Throughout we will always take the trait under study to be trait 1.

Trait specificity is then defined as importance for the trait of interest relative to importance across all fitness-relevant traits measured in the appropriate units (Figure 2**B**). We denote trait <u>sp</u>ecificity by 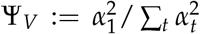 for variants and 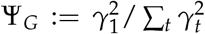 for genes. See Appendix A for more details. Ideally, association studies would prioritize genes based on trait importance, trait specificity, or some combination of the two.

### Burden tests prioritize trait-specific genes

To determine how LoF burden tests prioritize genes, we analyzed population genetics models of association studies (Appendix A). Our analysis revealed that LoF burden tests prioritize genes in part by their trait specificity, and not by importance (Figure 3**A**). We briefly outline the argument here.

**Figure 3:**
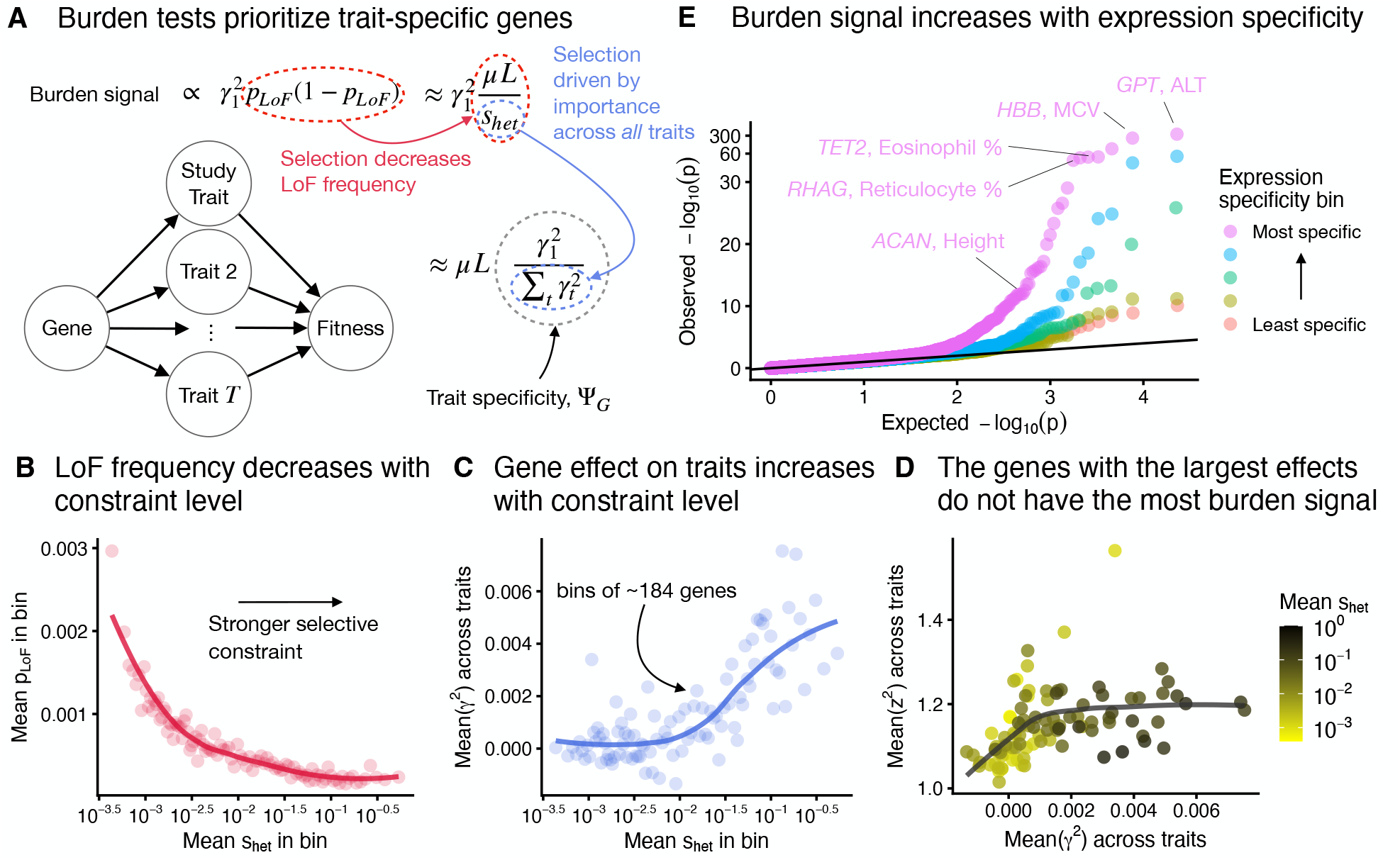
Burden tests prioritize trait-specific genes, not large-effect genes. **A**) Summary of our result that burden tests prioritize genes by trait specificity. µ is the per-site mutation rate, L the number of potential LoF positions, and s_het_ the strength of selection against heterozygous LoF carriers. **B**) Genes were binned by an estimate of s_het_ [33] with approximately 184 genes per bin. Aggregate LoF frequencies were then averaged across genes within each bin. The trend line was fit using LOESS. **C**) Similar to **B**, but averaging over an unbiased estimate of the mean of 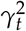 across traits. **D**) Genes were binned as in **B**, and the mean of squared z-scores, z^2^, across traits was plotted against the average of an unbiased estimate of the mean of 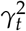 across traits. Points are colored by the mean s_het_ within the bin and the trend line was fit using LOESS. **E**) Quantile-quantile plot of LoF burden test p-values across 9 trait-tissue pairs. Genes were stratified for each trait-tissue pair based on the specificity of their expression to the trait-relevant tissue. The y-axis has been non-linearly transformed.

In an LoF burden test, the strength of association, *z*^2^, for a gene depends on both its trait importance,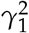, and the aggregate frequency of LoFs, *p*_*LoF*_, with the expected strength of association being proportional to 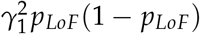.

Natural selection acts to keep LoFs rare: for sufficiently strong selection, *p*_*LoF*_(1 − *p*_*LoF*_) is proportional to *µL*/*s*_het_, where *µ* is the per-base mutation rate, *L* is the number of sites where an LoF could occur, and *s*_het_ is the strength of selection in heterozygotes [32]. As expected, there is a strong negative relationship between recently inferred estimates of *s*_het_ [33] and the average of *p*_*LoF*_ across genes within *s*_het_ bins (Figure 3**B**, Supplementary Figure S8).

Furthermore, many complex traits are thought to be under stabilizing selection [34–37]. Crucially, this predicts a connection between *s*_het_ and total trait effects. Specifically, 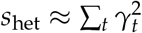, where 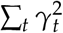 is the sum of trait importances across all fitness relevant traits measured in appropriate units (Appendix A). To test this prediction, we computed unbiased estimates of trait importance from LoF burden test results for 27 genetically uncorrelated traits (Methods). The average trait importance across these traits shows a strong positive relationship with *s*_het_ as predicted by our model (Figure 3**C**).

Combining these results, we see that the strength of association in LoF burden tests is proportional to 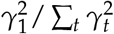 (Figure 3**A**). This is exactly our definition of Ψ_*G*_, the trait specificity of a gene.

A key implication of this result is that LoF burden tests do not prioritize genes based on trait importance. The most trait-important genes will often be the most constrained and have the smallest frequencies (and hence largest standard errors), an effect previously referred to as flattening [38, 39]. Indeed, Figure 3**D** shows that, for genes with sufficiently large effects, the strength of association 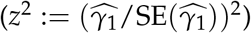 is completely decoupled from trait importance in the UKB LoF burden tests.

Instead, our theory predicts that LoF burden tests prioritize genes by their trait specificity, Ψ_*G*_. To confirm this prediction, we would ideally compare strength of association to an independent measure of Ψ_*G*_. It is difficult to directly estimate Ψ_*G*_ independently of our theory as it depends on the unknown true trait importances. Instead we use how specifically-expressed a gene is as a proxy. Our intuition is that genes expressed predominantly in a trait-relevant tissue are more likely to be trait specific than broadly-expressed genes.

To explore this, we selected 9 traits for which at least 40% of heritability was attributable to a single tissue in the ChIP Atlas [40]. For each trait, we focused on genes expressed in the top matched tissue and stratified them into quintiles of expression specificity, determined by their expression level in the focal tissue relative to the average across all the tissues we analyzed in the Human Protein Atlas (Methods) [41].

Using results from the LoF burden tests for these 9 trait-tissue pairs, we constructed a quantile-quantile plot (Figure 3**E**). Consistent with our intuition, we observed substantially stronger signals in the most specific expression bins. We also observed that many of the top hits are plausibly quite trait specific (Figure 3**E**). We found concordant results in a regression model that predicts LoF burden *z*^2^ from expression specificity even after controlling for differences in effect sizes (Supplementary Figure S9).

### GWAS prioritize trait-specific variants

We next turned to understanding how GWAS prioritizes genes. In contrast to LoF burden tests, GWAS are performed at the variant level, and so we consider what causes a variant to be ranked highly. Following the same argument as above reveals that the expected strength of association is proportional to 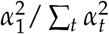, the trait importance of the variant for the trait under study relative to the total trait importance of the variant across all fitness-relevant traits. This is exactly Ψ_*V*_, the trait-specificity of the variant.

The fact that GWAS prioritizes trait-specific *variants* rather than *genes* has profound implications for understanding the differences between GWAS and LoF burden tests. In particular, variants can be trait specific in two ways (Figure 4**A**): they can either affect a trait-specific gene (e.g., variant 3 in Figure 4**A**) or affect a pleiotropic gene in a context-specific manner (e.g., variant 1 in Figure 4**A**). For example, context-specific variants might regulate expression only in trait-relevant cell types or at particular developmental time points, and thus have trait-specific effects even when acting on pleiotropic genes. In Appendix B, we develop a model formalizing the relationship between Ψ_*V*_, Ψ_*G*_, and context-specific expression.

**Figure 4:**
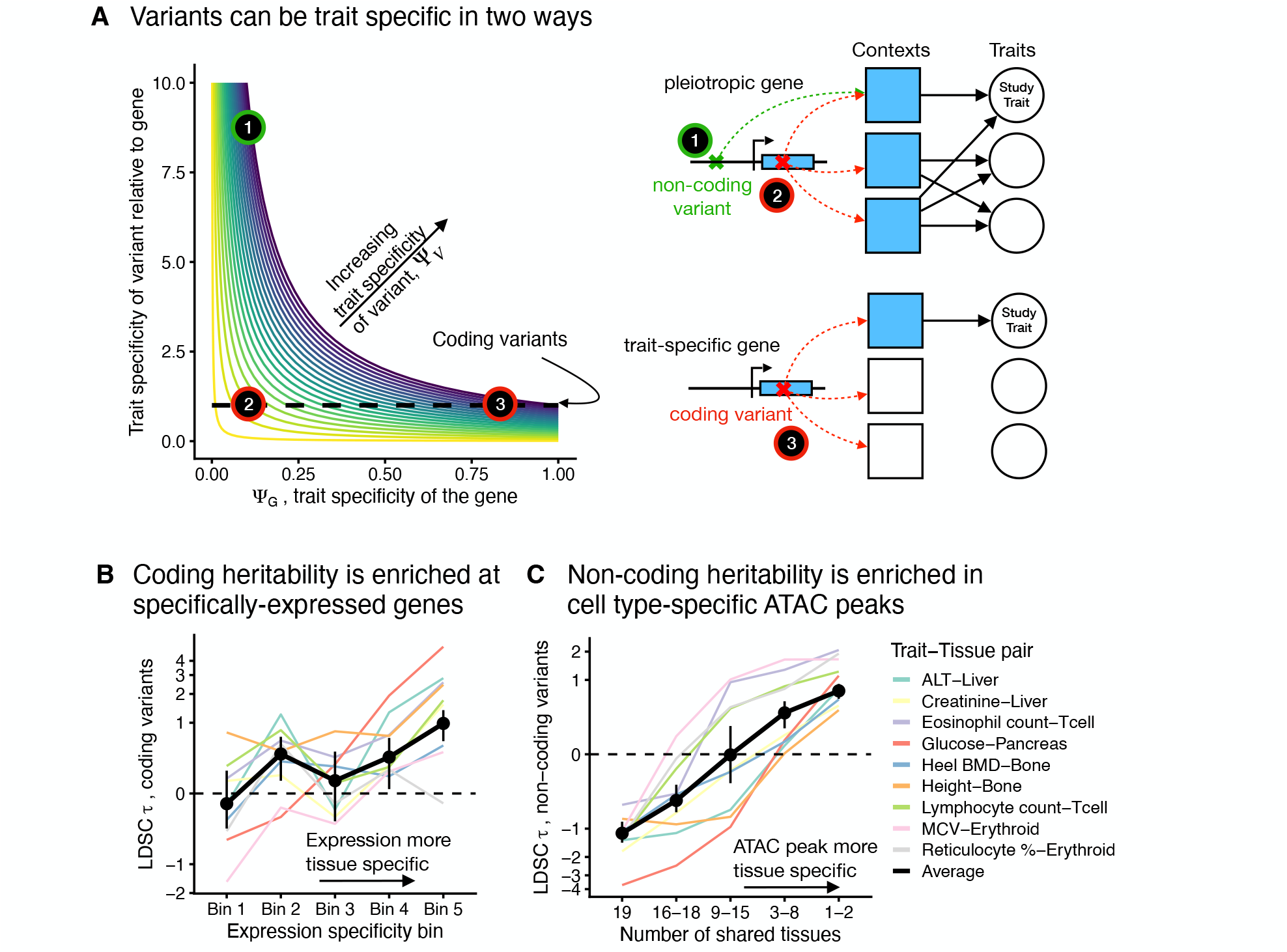
GWAS prioritize trait-specific variants. **A**) Schematic of what determines trait specificity for variants, Ψ_V_. Ψ_V_ is determined by both the trait specificity of the gene through which it acts and how trait specific it is relative to that gene. Three representative types of variants are highlighted with gene models. The green variant is non-coding, while the red variants are coding. Shaded contexts represent cellular contexts or cell types in which the gene affects traits. **B**) Heritability enrichment for coding variants as measured by S-LDSC τ as a function of expression specificity for 9 trait-tissue pairs. The inverse variance-weighted average of the results for the individual traits is in black. The y-axis has been non-linearly transformed. **C**) Heritability enrichment for non-coding variants in ATAC peaks as measured by S-LDSC τ as a function of how ATAC peak tissue specificity for 9 trait-tissue pairs. The inverse variance-weighted average of the results for the individual traits is in black. The y-axis has been non-linearly transformed.

To test our predictions, we considered the two ways that variants can be trait specific and used S-LDSC [6, 42] to quantify how heritability changes along these axes. The average heritability contributed by a set of variants is a proxy for how highly those variants would be prioritized by GWAS on average. We quantified effects on heritability by *τ* as reported by S-LDSC, which can be interpreted as how much a given annotation increases or decreases heritability.

First, we looked into whether the trait specificity of the gene on which a variant acts affects GWAS prioritization for variants with a given context specificity (moving along the horizontal axis of Figure 4**A**). To this end, we restricted our analyses to coding variants and again used the tissue specificity of each gene’s expression as a proxy for that gene’s Ψ_*G*_. Overall, variants acting on specifically-expressed genes are more likely to be prioritized highly by GWAS (Figure 4**B**, Supplementary Figure S9).

We next examined the impact of context specificity (moving along the vertical axis of Figure 4**A**). Here, we used non-coding variants, and we assumed that variants are more likely to have an effect in a tissue if they are in open chromatin in that tissue as determined by ATAC-seq [40]. We focused on the 9 trait-tissue pairs we analyzed in Figure 3**E**. For each variant in an ATAC peak in the top trait-relevant tissue, we then determined the number of additional tissues in which the variant is in open chromatin (Methods). We computed *τ* using S-LDSC as a function of this ATAC peak tissue specificity while controlling for the strength of the ATAC signal (Methods). Across all traits we see a strong trend of increasing heritability in more tissue-specific ATAC peaks (Figure 4**C**, Supplementary Figure S11).

Overall, our results show that LoF burden tests and GWAS both prioritize trait specificity, but prioritize different loci because LoF burden tests prioritize trait-specific *genes*, while GWAS prioritize trait-specific *variants*. Variants can be specific either by acting through trait-specific genes or by being context specific, and both of these axes contribute to a variant being prioritized by GWAS.

## Trait-irrelevant factors affect GWAS and LoF burden tests

Our modeling also revealed factors beyond trait specificity that affect which genes are prioritized by GWAS and LoF burden tests. These factors can cause one gene to be more highly prioritized than another for reasons that have nothing to do with their effects on any aspect of trait biology. LoF burden tests prioritize genes in part by the length of their coding sequence, and GWAS prioritize variants in part due to randomness in their frequencies caused by genetic drift.

### Gene length drives power in LoF burden tests

LoF burden tests aggregate all LoF variants within a gene (Figure 1**B**). As we derived above, this results in an expected strength of association that increases with *µL*, the average mutation rate times the number of potential LoF positions within a gene. Intuitively, if a gene has more potential LoFs, then the proportion of individuals that are LoF carriers will be larger, all else being equal, resulting in a higher aggregate LoF frequency and increased power.

Our model’s predictions about the impact of gene length are confirmed in the UKB LoF burden tests. We binned genes based on their expected number of unique LoFs, a measure of *µL* [43]. This measure is strongly correlated with coding sequence (CDS) length (Supplementary Figure S12), so we refer to this as “gene length”. We computed unbiased estimates of the average squared effect size of the genes, finding little association between gene length and total effect size (Figure 5**A**). Meanwhile, longer genes have considerably smaller standard errors on average (Figure 5**B**). Together, this results in the strength of association (*z*^2^) correlating strongly with gene length (Figure 5**C**), even though longer genes are generally not more trait important.

**Figure 5:**
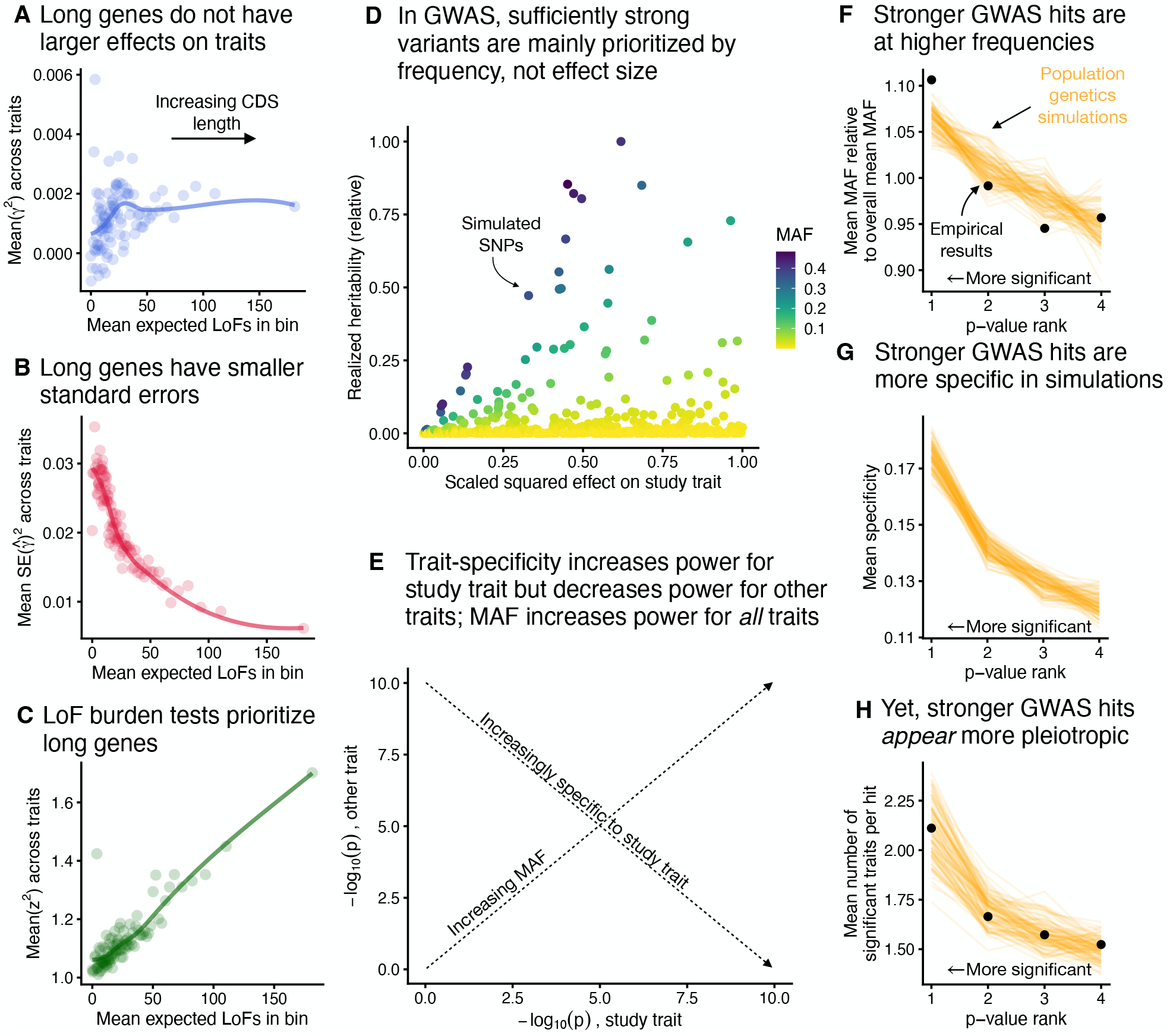
Trait-irrelevant factors drive prioritization in association studies. **A**) Average of an unbiased estimate of the squared trait importance, γ^2^, across 27 genetically uncorrelated traits, averaged within bins of approximately 184 genes binned by expected number of unique LoFs [43]. The trend line was fit using LOESS. This analysis was repeated for **B**) the average of the squared LoF burden test standard errors within each bin, and **C**) the average LoF burden test z^2^ across traits within each bin. **D**) Simulations of realized heritability for individual variants with varying trait importances, scaled by the maximum simulated realized heritability. **E**) Schematic of the effects of minor allele frequency (MAF) and trait specificity on GWAS p-values. **F**) The relationship between MAF and p-value rank for simulations and real data across genetically uncorrelated traits. Genome-wide significant hits were binned by p-value, and the mean MAF within each bin was compared to the overall mean MAF across all hits. Black points are results from UKB GWAS, and orange lines are simulations. This analysis was repeated for **G**) the mean trait specificity within each bin and **H**) the mean number of traits for which each hit was genome-wide significant. Panel **G** contains only simulations as the trait specificities for the UKB GWAS results are unknown.

### Random genetic drift drives power in GWAS

We showed above that the expected strength of association in GWAS is proportional to a variant’s trait specificity, Ψ_*V*_. This is true on average, but there is considerable variation around this expectation. In well-powered GWAS, variants are ranked by 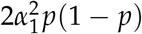, where *p* is the variant allele frequency (Appendix A). We refer to 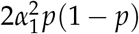 as a variant’s *realized heritability*. Under our modeling assumptions, the expected value of *p*(1 − *p*) is 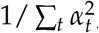, resulting in trait-specific variants being ranked more highly *on average*. Yet, random genetic drift causes variant frequencies to be spread widely around their expected values (Supplementary Figure S13).

In LoF burden tests this effect is largely ameliorated by the aggregation of variants, which averages out the stochasticity in the frequencies of individual LoFs (Appendix A). But GWAS consider variants one at a time, causing this stochasticity to play a large role in gene prioritization.

In Figure 5**D**, we plot simulations of realized heritability under our model (Methods). The ranking of variants in terms of realized heritability is largely random with respect to trait importance for sufficiently trait-important variants. This randomness is driven by differences in minor allele frequency (MAF) caused entirely by genetic drift.

This randomness in MAF explains an apparent contradiction between our finding that GWAS prioritize trait-specific variants and previous studies that report GWAS hits appearing to be surprisingly pleiotropic [44–46]. Consider performing GWAS on two traits (Figure 5**E**). If a variant is trait specific for one trait, then by definition it cannot be trait specific for the other trait. In the absence of other forces, this results in a negative relationship between the strength of association for the two traits. In contrast, if a variant has high MAF, then the GWAS for both traits will be well-powered. All else being equal, randomness in MAF results in a positive relationship between the strength of association for the two traits. Therefore, variants that are highly ranked in one GWAS will be enriched for variants that are trait specific (and hence less likely to be hits for the other trait) but also variants that have high MAF (and hence are more likely to be hits for the other trait). This explains the supposed contradiction: the top hits for one trait are not actually more pleiotropic on average than other variants, they are simply at higher MAFs and hence better powered across all traits.

To see if this prediction of our model is corroborated by the UKB GWAS, we compared properties of GWAS hits taken from 27 genetically uncorrelated traits to properties of GWAS hits simulated under our model (Methods). In both cases, we considered all variants that passed the genome-wide significant threshold as hits, and then partitioned hits into four bins based on their p-values, with the strongest hits being in bin 1 and the weakest (but still genome-wide significant) hits being in bin 4.

Our model recapitulates the behavior of the UKB GWAS hits. As predicted by Figure 5**E**, the strongest GWAS hits are at higher than average frequencies in both our model and the UKB GWAS hits (Figure 5**F**). As mentioned above, trait specificity is difficult to directly measure, so we cannot assess the trait specificity of the real GWAS hits, but in our simulations, GWAS does indeed prioritize trait-specific variants (Figure 5**G**). Finally, consistent with previous studies [44–46], we find that the top GWAS hits for one trait are hits for other traits more often than weaker GWAS hits (Figure 5**H**). We reiterate that on average these hits are in fact *more* trait-specific despite being GWAS hits for more traits, and this discrepancy is caused by their higher than expected MAF. The precise details of our simulation model have a quantitative, but not qualitative effect on these results (Methods; Supplementary Figures S14–S17)

### Approaches for estimating trait importance

We began by proposing that it could be desirable to prioritize genes either by trait importance or trait specificity. Yet, so far we have shown that when ranking by p-value neither LoF burden tests nor GWAS rank genes by their trait importance. We wanted to see if there was some way to use GWAS or LoF burden test results to prioritize genes in a way that is correlated with trait importance.

In this section, it will be helpful to consider a simplified model where each variant has an effect *β* on some gene that has effect *γ* on the trait, so that the overall effect of the variant on the trait is *α* = *βγ* (similar to the models in [47]). This assumption is for ease of exposition: in reality, we have found that *α* often depends on *β* non-linearly [48], but this does not qualitatively affect our results.

Throughout, we have focused on prioritizing genes based on p-value or strength of association. It is then natural to ask if ranking genes on some other summary of the association tests would better prioritize important genes (e.g., the unbiased estimates of trait importance 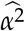 for GWAS or 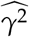 for LoF burden tests). For GWAS, we immediately see that this is not possible without additional information: the relationship between any estimate of *α* and trait importance will depend on the unknown value of *β*.

In principle, if an LoF burden test is infinitely well powered, then ordering genes by 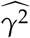 would prioritize genes based on trait importance. At current sample sizes, however, the estimated 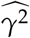 are noisy enough that the top genes will contain many false positives. For example, among the 10 genes with the largest 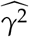 for standing height, 4 are consistent with actually having no effect on height (all Bonferroni adjusted p-values > 0.62). Furthermore, if LoFs in a given gene are extremely deleterious, then LoF burden tests may never be well powered no matter the sample size, resulting in false negatives.

Estimating trait importance is most difficult for the most important genes, a phenomenon called *flattening* [19, 38, *39]*. Flattening refers to the expected strength of association (equivalently the expected contribution to heritability) first increasing as (βγ)^2^ increases, but then becoming uncoupled from (*βγ*)^2^ for sufficiently large (*βγ*)^2^ (Figure 6**A**; Appendix C). This decoupling causes association studies to be incapable of prioritizing by trait importance, which we see in LoF burden tests (Figure 6**B**), where contributions to heritability are completely uncoupled from *s*_het_, which we use as a proxy for trait importance based on Figure 3**B**.

**Figure 6:**
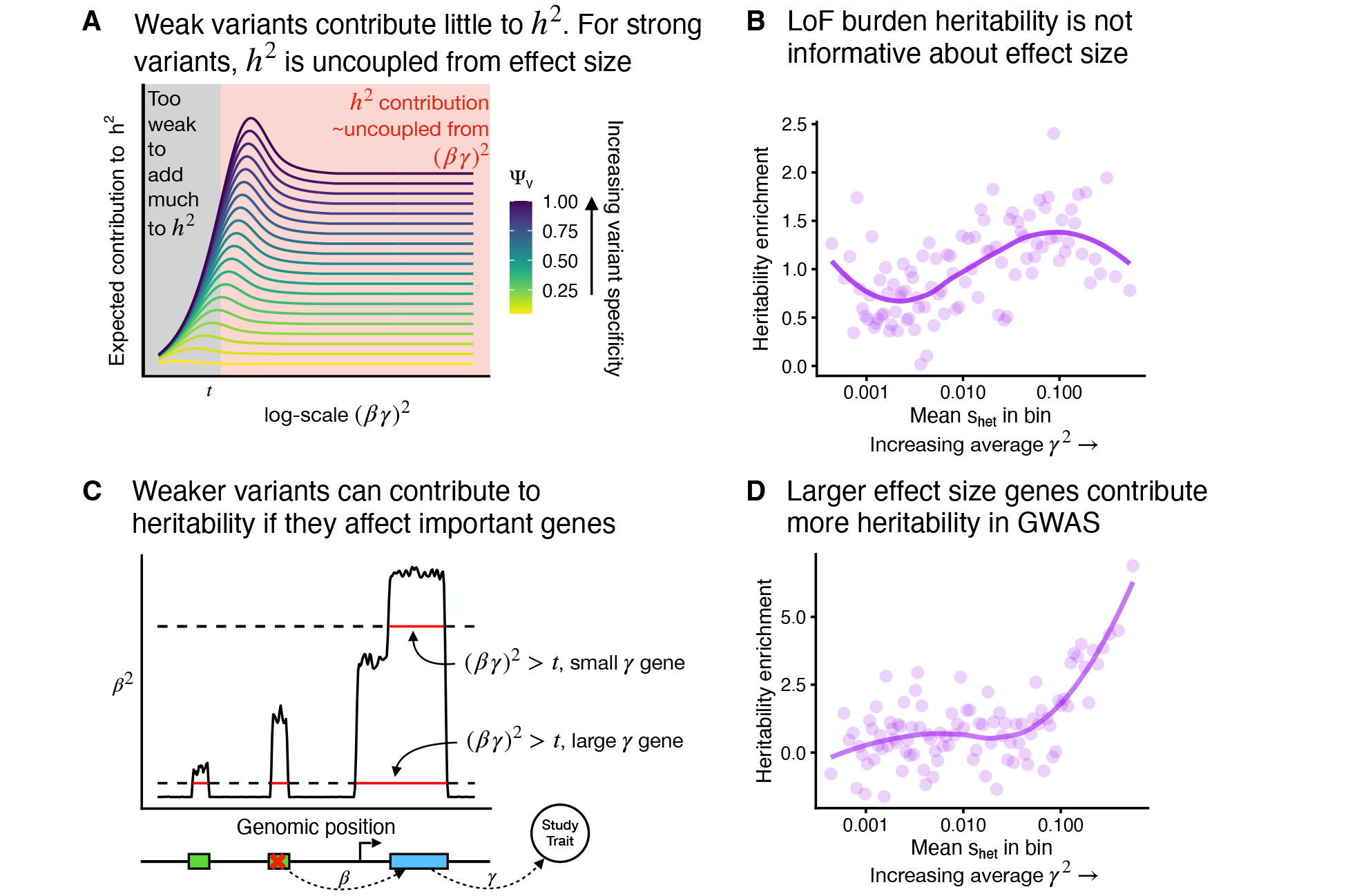
Estimating trait importance by combining information across different variant types. **A**) Theoretical expected contributions to heritability, h^2^, as a function of the total effect of a variant on a trait α^2^ = (βγ)^2^. Colored lines are variants with different trait specificities. These functions can be approximately divided into a regime where variants contribute vary little to heritability (black) or their contribution depends very little on (βγ)^2^ (red). **B**) Enrichment of LoF burden test heritability for genes binned by selective constraint s_het_. The plotted heritability enrichment is a normalized inverse variance-weighted average of heritability enrichments across 27 genetically uncorrelated traits (Methods). Each bin contains approximately 184 genes. The trend line was fit using LOESS. **C**) Schematic of how a variant’s contribution to heritability depends on the γ^2^ of the gene through which it act. Green boxes represent cis regulatory regions. **D**) Similar to panel **B**, but estimated using GWAS results instead of LoF burden test results. Per-trait estimates were obtained using AMM [49], and we plot a normalized inverse variance-weighted average across the same traits as in Panel **B** (Methods).

Yet, flattening does not affect all variants in the same way. For simplicity we can imagine that variants either contribute minimally to heritability, or they contribute an amount independent of their importance (Figure 6**A**). Now, imagine two genes: one has a large effect on the trait (large *γ*), and one has a small effect (small *γ*). Even variants that weakly perturb the large *γ* gene will have large enough (*βγ*)^2^ to contribute to heritability, whereas for the small *γ* gene, only variants with very large *β* will contribute to heritability (Figure 6**C**). Each individual variant will experience flattening, but collectively there will be more variants that contribute to heritabilty for more trait-important genes, all else being equal. As a result, we expect the total heritability contributed by variants acting on a given gene to correlate with that gene’s trait importance.

To test this prediction of our model, we used AMM [49], a method that estimates the heritability of variants acting via a given set of genes using GWAS summary statistics (Methods). We found that compared to LoF burden heritability, this measure of total heritability better tracks *s*_het_ and hence trait importance (Figure 6**D**). The results of this analysis do not rely on the specific details of AMM: we expect similar results whenever aggregating signals across variants with different *β* (e.g., Supplementary Figures S18 and S19; Methods; and [50]).

## Discussion

It is often assumed that GWAS and burden tests converge on similar gene sets [18, 51, 52]. Indeed, some genes are implicated by both approaches, such as *LDLR* for low-density lipoprotein levels [19, 53]. Generally, GWAS loci are enriched near burden genes and, conversely, burden genes — as well as genes identified in familial studies of Mendelian counterparts of the same traits — are enriched within GWAS loci [16, 54].

Here, we find that, despite this overall concordance, LoF burden tests and GWAS rank genes differently, resulting in limited overlap among the top genes identified by each approach. Our analysis shows that LoF burden tests prioritize long, trait-specific genes, while GWAS prioritize genes near trait-specific variants that have drifted to unexpectedly high frequencies. Because context-specific variants can be trait-specific even if they act on pleiotropic genes, GWAS can find trait-relevant, pleiotropic genes that would be missed by LoF burden tests.

These findings have significant implications for interpreting GWAS and LoF burden tests and their applications. They help explain why burden tests often appear less polygenic than GWAS and tend to prioritize genes that are seemingly more directly related to trait biology [19]. GWAS should, in principle, capture all trait-specific genes identified by LoF burden tests, but also identify highly pleiotropic, selectively constrained, trait-relevant genes.

The fact that we find numerous examples of GWAS loci with essentially no LoF burden signal suggests that such highly pleiotropic genes are major drivers of complex traits. We hypothesize that some of these genes have developmental roles and that GWAS heritability is partly driven by context-specific variation that perturbs developmental trajectories in a trait-specific manner [55].

While both study designs identify sufficiently trait-important genes, neither directly ranks genes by trait importance. LoF burden tests estimate trait importance, but selection causes estimation noise to increase with gene effect size, making rankings by significance nearly independent of trait importance. Gene length is also a major confounder. Although larger sample sizes will help reduce noise, we anticipate that Bayesian frameworks using priors based on gene features, such as our recent approach to estimating *s*_het_ [33], could be particularly effective for improving the accuracy of burden tests.

In GWAS, genetic drift makes the p-values of individual variants essentially arbitrary as long as the variants are sufficiently trait specific and important. This makes variant-level ranking of GWAS loci inefficient for identifying top genes. Instead, genes can be prioritized by trait importance using non-standard GWAS approaches that aggregate signals across multiple variants (e.g., [49, 50, 56, 57]), motivating further development of such methods.

Our findings also explain why GWAS results are highly effective for identifying trait-relevant tissues and cell types using approaches like S-LDSC [6]. Variants that are only active in trait-relevant cell types are much more likely to be trait specific, and thus contribute more to heritability. This is not necessarily because such variants have larger effect sizes, but rather that for a given effect on the trait they are at higher frequencies on average.

The question of how genes should ideally be prioritized is surprisingly understudied. Here we propose ranking genes based on either trait importance or trait specificity. Both concepts capture different aspects of what it means for a gene to be “relevant” for a trait. Is a gene that has only a modest effect on a trait, but affects no other traits more or less relevant than a gene that has massive effects across a whole suite of unrelated traits?

Which prioritization is more useful will likely depend on the downstream context. For example, trait-specific genes may be better drug targets due to reduced side effects, perhaps explaining why LoF burden evidence is more predictive of drug trial success than GWAS evidence [58]. Yet, if pleiotropic genes can be targeted in a context-specific way, perhaps prioritizing genes by trait importance may identify the most impactful therapeutic targets. Additionally, the effects of pleiotropic genes in knockout experimental systems may differ fundamentally from the phenotypic consequences of regulatory variants identified in GWAS.

The fact that LoF burden tests and GWAS prioritize different genes is a blessing — both are useful and both reveal different aspects of trait biology. However, it is important to understand what genes they prioritize and why. Our results make clear that both association study designs will be important in future efforts to map the genetic underpinnings of complex traits.

## Methods

### GWAS summary statistics

GWAS summary statistics for 305 continuous traits were downloaded from the Neale Lab (http://www.nealelab.is/uk-biobank/, version 3). These regressions were run on inverse rank Normal transformed phenotypes in a subset of the UKB consisting of approximately 360,000 individuals and included `age`, `age`^2^, `inferred_sex`, `age` × `inferred_sex`, `age`^2^ × `inferred_sex`, and principal components 1-20 as covariates. We used 5 `×` 10^−8^ as the threshold for genome-wide significance unless otherwise stated.

### LoF Burden test summary statistics

Summary statistics for 292 LoF burden tests were downloaded from Backman *et al*. [16]. 209 traits overlapped with traits for which we had GWAS summary data (Supplementary Table 1). Burden genotypes were calculated for each individual by assigning a homozygous reference genotype to individuals homozygous for the reference allele for all considered LoF variants and assigning a heterozygous genotype to all other individuals. Burden tests were run using REGENIE [59], on inverse rank Normal transformed phenotypes. For all analyses, we used the result of the burden test with mask M1, which only includes variants that are predicted as being LoFs using the most stringent filtering criteria and an allele frequency upper bound of 1%. We used a per-trait genome-wide significance threshold of 2.7 × 10^−6^, derived by applying a Bonferroni correction to a significance threshold of 0.05 for testing approximately 18,000 genes per trait.

### A subset of genetically uncorrelated traits

The set of 209 quantitative traits included some that were highly correlated, such as *sitting height* and *standing height*. For certain analyses, we selected a subset of 27 traits that were not highly correlated by intersecting the 209 traits with those analyzed by Mostafavi *et al*. [47] (Supplementary Table 1). Briefly, the trait list was pruned to ensure that all pairwise genetic correlations, as reported by the Neale lab, were below 0.5, prioritizing traits with higher heritability. Biomarkers were excluded from this subset because their genetic correlations with other traits were not provided by the Neale lab.

### Defining GWAS loci

For a systematic comparison of discoveries between GWAS and burden tests (shown in Figures 1**C** and 1**D**), we grouped GWAS variants into large, non-overlapping genomic loci. This approach avoids multiple counting of the same GWAS genes, as nearby hits within a locus may map to the same gene, and it provides a conservative estimate of the overlap between GWAS and burden test results as described below.

We focused on 151 quantitative traits with at least one burden test hit and one GWAS hit. For each trait, we analyzed the set of LD-clumped hits (*p* < 5 × 10^−8^, clumping *r*^2^ < 0.1) from 8,136,100 filtered SNPs provided by Mostafavi *et al*. [47].

For each trait, we began the grouping procedure with the most significant hit and iteratively processed all hits until they were assigned to a locus. For each hit, we included all independent hits with larger p-values (lower significance) within 1Mb to form a locus. The locus size was then expanded to ensure that no other hit was within 1Mb of any variant already included in the locus. After completing one locus, we moved on to the next most significant hit that had not yet been assigned to any locus. Finally, we assigned overlapping genes to each locus, focusing on the 18,524 protein-coding genes analyzed in the LoF burden test.

In Figure 1**D**, we plot the p-value of the most significant GWAS variant within each locus on the x-axis and the p-value of the most significant gene within the same locus from the burden test on the y-axis.

In Figure 1**C**, we include only the top GWAS loci to match the statistical power of the burden test for gene discovery. We illustrate our procedure with the example of standing height. The LoF burden test for standing height identified 82 significant genes (*p* < 2.7 × 10^−6^, to account for the 18,524 genes tested). The GWAS analysis identified 3,374 nearly independent hits. Following the grouping procedure outlined above, these hits were consolidated into 382 loci (median size 3.2 Mb). We ranked these loci by the minimum p-value within each locus. Starting with the top-ranked locus, we iteratively added GWAS loci until we selected 82 genes. From each locus, we selected all genes that were significant in the LoF burden test. If no such genes existed, we selected the gene with the smallest burden test p-value.

This procedure ensures that our analysis of the overlap between burden test and GWAS discoveries is conservative. The overestimation arises first from prioritizing genes based on burden test p-values and second from using large GWAS loci, which may contain more than one causal gene, thereby increasing the likelihood of overlap with burden test results

### Comparing GWAS and LoF burden tests at the LD block level

To avoid exacerbating dissimilarities between LoF burden tests and GWAS caused by mislocalization of GWAS signals, we also performed analyses at the LD block level. We downloaded bed files containing the coordinates of approximately independent LD blocks from [60]. For each trait we computed the minimum GWAS p-value of variants within each block and compared that to the minimum LoF burden test p-value for all genes that overlapped any part of that block. In a small number of cases, the smallest LoF burden test p-value in two adjacent blocks would be the same because a single highly significant gene overlapped both blocks. This generally reduced the correlation between the minimum p-values of GWAS and LoF burden tests, and so we dropped all such blocks to be conservative.

### Association study model

We combined population genetics and statistical genetics models to understand how natural selection affects variants based on their trait specificity and trait importance. Our model assumes that traits are under stabilizing selection based on prevailing hypotheses [34, 36, 37], and uses standard population genetics theory [38, 61–64]. The details of our model are outlined in the Appendices.

### Unbiased estimates of trait importance

In several analyses we require estimates of trait importance, either *α*^2^ from GWAS or *γ*^2^ from LoF burden tests. The details in both cases are identical, so here we describe *γ*^2^. The naïve estimator of squaring the LoF burden test estimated effect size, 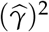 is biased. Worse, this bias is anticorrelated with the frequency of the variant, which results in spurious correlations between the biased estimates and various gene properties such as *s*_het_.

To derive an unbiased estimator, we appeal to standard statistical genetics theory [65] to assume that LoF burden estimates are approximately Normally distributed about their true values with noise dependent on their standard errors. In particular, for a gene with standard error *s* and effect size estimate 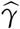, we have that 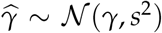 approximately. This approximation is widely-used for for GWAS and was recently confirmed to be accurate for LoF burden tests [48]. It is then a routine calculation to check that 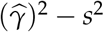 is an unbiased estimator of *γ*^2^.

### LoF burden summary statistics as a function of *s*_**het**_

When comparing LoF burden summary statistics (standard errors, *z*^2^, and unbiased estimates of *γ*^2^,) to *s*_het_, we used *s*_het_ values inferred in [33] and downloaded from [66]. We binned genes by *s*_het_ into 100 bins, each with approximately 184 genes. Within each bin we averaged the respective summary statistics (e.g., unbiased estimate of *γ*^2^) across traits and genes. To make sure that our results were not driven by redundant traits, we used our 27 genetically uncorrelated traits for these analyses. For heritability enrichment (Figure 6**B**), we used the fact that heritability should be proportional to *z*^*2*^−1 (Appendix A). Within each bin of genes we then computed the average *z*^*2*^−1 in that bin relative to the average of *z*^*2*^−1 across all genes for each trait. This produced a trait-level enrichment for each bin, and by using the empirical standard deviation of the relative *z*^*2*^−1 within the bin, we could also obtain an empirical standard error for the enrichment. We then obtained an overall enrichment for each bin, by taking an inverse-variance weighted average across traits. After this averaging, the mean heritability enrichment across genes need not be one. As such, we renormalized the estimates to average to one.

### ATAC peak specificity

We downloaded all ATAC-seq files from ChIP-Atlas [40] that contained more than 5,000,000 mapped reads and identified at least 5,000 peaks. Across all files, overlapping peaks were combined using bedtools merge [67]. This yielded a total of 2,131,526 peaks. Samples other than blood samples were grouped into 17 tissues based on their annotations in ChIP-Atlas. Namely, Adipocyte (146 samples), Bone (190 samples), Breast (815 samples), Cardiovascular (559 samples), Digestive (417 samples), Epidermis (661 samples), Gonad (138 samples), Kidney (375 samples), Liver (191 samples), Lung (1679 samples), Muscle (118 samples), Neural (1349 samples), Pancreas (322 samples), Placenta (48 samples), Pluripotent (1895 samples), Prostate (312 samples), and Uterus (255 samples). Additionally, samples with any of the following annotations were categorized as Tcell (1356 samples): CD4-Positive T-Lymphocytes, CD4+ T cells, CD8-Positive T-Lymphocytes, CD8+ T cells, Fetal naive T cells, Gamma-delta T cell, Naive T cells, T cells, CAR-T cells, Tfh, Th0 cells, Th17 Cells, Th1 Cells, Th2 Cells, Th9 Cells, or T-Lymphocytes. Samples with any of the following annotations were categorized as Erythroid (102 samples): Erythroid progenitors, Erythroid Cells, or Erythroblasts. Ultimately this resulted in 19 tissues or cell type categories.

A peak was considered to be present in a tissue if more than 5 percent of samples contained the peak. In downstream analyses we used both “number of shared tissues” and “peak intensity”. We calculated the number of shared tissues by considering all peaks in the relevant tissue for a given trait (e.g., Bone for Height) and then counting the number of tissues in which that peak was present. In particular, we only consider peaks that are present in the relevant tissue. We calculated “peak intensity” as the fraction of samples within the focal tissue that contain the peak.

### Gene expression specificity

We compiled estimates of gene expression in 17 tissue/cell types, which were intended to overlap with the categorization of ATAC-seq peaks when possible. All tissues that were ultimately matched to traits (see below) were included in both our ATAC-seq tissues and our expression tissues, but there are some differences between the remaining tissues. Average gene expression TPM of the following tissues were downloaded and extracted from the Human Protein Atlas [41] tissue gene data (`rna_tissue_hpa.tsv.zip`): adipose tissue, breast, heart muscle, colon, skin, ovary, kidney, liver, lung, skeletal muscle, amygdala, pancreas, placenta, and prostate. Average gene expression TPM of the following cell types were downloaded and extracted from the Human Protein Atlas single cell type data (`rna_single_cell_type.tsv.zip`): Erythroid cells and T cells. Average gene expression TPM of human bone samples was downloaded from GEO [68] accession GSE106292 [69, 70].

In each tissue, genes with more than 10 TPM were considered to be “expressed”. We then restricted our analyses to genes expressed in the trait-relevant tissue. We computed an expression specificity score by taking the expression level in TPM in the trait-relevant tissue divided by the sum of expression levels across all 17 tissues. This provided an expression specificity score for every gene expressed in the trait-relevant cell type. For analyses involving expression specificity bins, we took all of these expression specificity scores across all 9 trait-tissue pairs, computed quintiles, and then assigned each gene for a given trait-tissue pair to its quintile.

### Linking traits to tissues

To identify which tissue (or cell type) is predominantly associated with a given trait, we ran S-LDSC [6, 42] to partition the heritability of all of our traits that had estimated heritability > 0.04. We used annotations for 19 tissues and cell types constructed from our ATAC-seq analysis described above, along with the LDSC baseline v1.1 covariates. Our aim was to identify trait-tissue pairs where heritability could clearly be explained by one tissue as opposed to multiple tissues. As such, we only retained traits that had a tissue with an LDSC *τ* with a *z*-score > 4.5 and had>40% of their heritability explained by variants in ATAC-seq peaks of the corresponding tissue. If more than one trait was assigned to the same tissue, we only kept genetically uncorrelated traits (*r*^2^ < 0.04). This resulted in 9 trait-tissue pairs (Supplementary Table 1): Mean corpuscular volume (30040_irnt) → Erythroid, Reticulocyte percentage (30240_irnt) → Erythroid, Eosinophil percentage (30210_irnt) → T cell, Lymphocyte count (30120_irnt) → T cell, Standing height (50_irnt) → Bone, Heel bone mineral density (3148_irnt) → Bone, Glucose (30740_irnt) → Pancreas, Creatinine (30700_irnt) → Liver, and Alanine aminotransferase (30620_irnt) → Liver.

### Estimating the effect of gene expression specificity on LoF burden prioritization

For each of the 9 trait-tissue pairs described above, we performed a linear regression of the burden *z*^2^ for all genes expressed in the top tissue on the genes’ expression specificity, binned into quintiles as described earlier. We included the unbiased estimates of the genes’ trait importance (defined above) as a covariate. For each specificity bin, we calculated an inverse-variance weighted average of the regression coefficients across all 9 traits, with standard errors computed as the square root of the reciprocal of the total weight. The results, shown in Supplementary Figure S9, demonstrate that the burden test prioritization of specifically expressed genes in Figure 3**E** is not driven by differences in the importance of genes across specificity bins.

### S-LDSC analysis using tissue-specific ATAC-seq peaks

For each trait-tissue pair, we ran S-LDSC [6, 42] to estimate the heritability enrichment of tissue-specific ATAC-seq peaks. To this end, we categorized ATAC-seq peaks present in each tissue into 5 bins based on their presence in other tissues: present in 1-2 tissues, present in 3-8 tissues, present in 9-15 tissues, present in 16-18 tissues, and present in all 19 tissues. Also, we categorized ATAC-seq peaks present in each tissue into 5 bins based on their intensity. The size of these bins were set to match the sizes of the tissue-specificity-based bins. We included the annotations based on ATAC peak tissue specificity and peak intensity bins with the LDSC baseline v1.1 model and used S-LDSC v.1.0.1 on HapMap3 SNPs [71].

### S-LDSC analysis using coding variants

We downloaded the variant annotation file (variants.tsv.bgz) from the Neale lab website ((http://www.nealelab.is/uk-biobank/). We used the consequence information in the file, which corresponds to Ensembl Variant Effect Predictor (VEP) version 85 [72], for annotating variants. Specifically, we classified variants as being coding if their most severe consequence was any of:

- `splice_donor_5th_base_variant`
- `missense_variant`
- `splice_region_variant`
- `splice_acceptor_variant`
- `splice_donor_variant`
- `splice_donor_region_variant`
- `stop_gained`
- `start_lost`
- `stop_lost`
- `frameshift_variant`
- `inframe_insertion`
- `protein_altering_variant`

For each trait–tissue pair, we ran S-LDSC [6, 42] to estimate the heritability enrichment of coding variants as a function of expression-specificity. We included expression specificity bin (as defined above) as an annotation in the S-LDSC model. We also categorized genes into 5 equally sized bins based on their expression level in the tissue of interest, and all the coding variants were categorized into one of these 5 bins based on the expression level of the corresponding genes. These annotations were also included in the S-LDSC model. In addition, we used the covariates in the baseline v1.1 model and restricted our analysis to HapMap3 SNPs [71]. All analyses were run with S-LDSC v.1.0.1.

### LoF burden summary statistics as a function of *µL*

Analyses comparing LoF burden summary statistics to *µL* were performed analogously to the analyses comparing the summary statistics to *s*_het_. As a proxy for *µL*, we downloaded the expected number of segregating LoFs for each gene as calculated in gnomAD v2 [43] from [66]. To show that *µL* is essentially driven by CDS length, we downloaded CDS lengths for MANE select canonical transcripts (genome build GRCh38) from ensmbl [73] and correlated them with the expected number of segregating LoFs from gnomAD [43] (Supplementary Figure S12).

### Computing variant frequency distributions as a function of *s*_**het**_

To simulate under our model, we required the distribution of allele frequencies for a given selection coefficient. We assumed a stabilizing selection model which is approximately equivalent to homozygotes having a relative fitness of 1 and heterozygotes having a fitness of 1 − *s*_het_ [38,61–63]. We used `fastDTWF` [74] to compute likelihoods under this model. We assumed an equilibrium population of 20,000 diploids, and computed allele frequency distributions along a grid of 50 *s*_het_ values from 10^−7^ to 0.05 evenly spaced on the log scale. We used 1.25 × 10^−8^ as the per-generation mutation rate. We considered a model where the ancestral allele is known by using `fastDTWF`’s `no_fix=True` option. Additionally, `fastDTWF` has two parameters that control the accuracy of its approximation. Based on the recommendations of [74], we set `dtwf_tv_sd` to 0.1 and `dtwf_row_eps` to 10^−8^.

### Simulating realized heritability

To generate Figure 5**B**, we simulated 50,000 unlinked variants from our stabilizing selection model. We considered 1, 000 values of *s*_het_ log-uniformly spaced between 10^−7^ and 2.3 × 10^−4^. For each value of *s*_het_, we then simulated 50 variants by drawing 50 allele frequencies from the allele frequency distributions we computed as described above. To model the slight differences between population and GWAS sample allele frequencies, we then drew a GWAS sample allele count for each variant as Binomial(600,000, *f*) random variable, where *f* was the population frequency, and 600,000 was chosen to match the roughly 300,000 diploids in the UKB. These allele counts were then normalized to obtain GWAS sample allele frequencies, 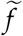. For this simulation, we assumed that all variants have the same trait specificity. This makes *α*^2^ on the focal trait proportional to *s*_het_, so we set the realized heritability to 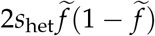 and normalized all results relative to the maximum simulated realized heritability. Likewise, effect sizes were reported relative to the maximum simulated effect size.

### Computing pleiotropy of GWAS hits

To investigate the pleiotropy of top versus weak GWAS hits, we considered all of the 27 uncorrelated traits that had at least 100 GWAS hits, leaving 18 traits. For each trait, we grouped the hits into four quartiles based on variant p-values, with quartile 1 containing the most statistically significant hits and quartile 4 the least. For each hit, we calculated the number of traits (out of 18) in which the variant was a hit and computed the mean values within each quartile.

### Simulating pleiotropy of GWAS hits

To simulate the effects of genetic drift on the apparent pleiotropy of GWAS hits, we simulated GWAS summary statistics. To match the real data described above, we considered 18 traits, and simulated effect sizes for 10 million not necessarily segregating positions. We simulated the effect sizes independently for each position, and drew the vector of squared effect sizes for variant *j*,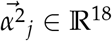 as

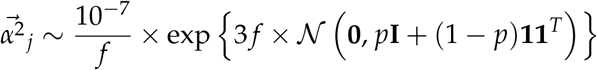

where the exponentiation is performed element-wise and *f* and *p* are parameters that affect the range of different total effect sizes,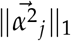, and distribution of trait specificities.

We then assumed that the strength of selection against the variant was 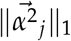. We obtained the minor allele frequency for each variant by drawing from the variant frequency distribution with the closest *s*_het_, computed as described above.

Finally, we simulated a GWAS by assuming that the observed association statistic for each trait was independently Normally distributed about its true value. For example, for trait *k* and the variant at position *j* we have:

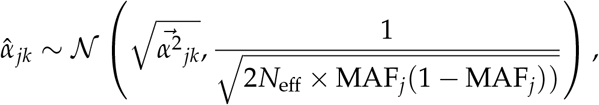

where *N*_eff_ is a scaling factor that captures both the the amount of environmental noise conributing to the trait as well as the sample size. We converted these to p-values by taking 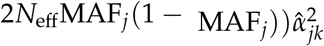 as a squared z-score, which is chi-squared distributed with one degree of freedom under the null. We considered a variant to be a genome-wide significant hit if its p-value was smaller than a parameter, *t*.

This simulation approach has four free parameters. In the main text we use *f* = 0.33, *p* = 0.5, *N*_eff_ = 10000000 and *t* = 10^−5^. While these parameters are related to standard GWAS parameters (e.g., the GWAS sample size or genome-wide significance threshold), the exact quantitative relationship should not be over-analyzed. For example, we assume that the strength of selection is exactly 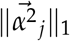 If instead there was some scaling factor, that could be absorbed into *N*_eff_. Similarly, there is a qualitative inverse relationship between the effects of *t* and *N*_eff_ (e.g., lower *t* has a similar effect to increasing *N*_eff_), making the exact setting of either parameter somewhat arbitrary. We chose the values we used here to roughly match the distribution of selection coefficients inferred from real GWAS data [75], as well as the observed patterns of MAF and pleiotropy in the UKB GWAS results. In Supplementary Figures S14, S15, S16, and S17, we vary each of *N*_eff_, *t, p*, and *f* respectively while holding the others fixed to show that our qualitative results are not sensitive to the particular simulation parameters we chose.

### AMM analysis

We ran AMM [49] to estimate heritability enrichments for gene sets, following the workflow described at (https://github.com/danjweiner/AMM21 commit 524c620). We binned genes into 100 approximately equally sized bins based on *s*_het_ as described above and used these bins as our gene sets. AMM requires an estimate of the probability that a SNP is acting via the closest gene, second closest gene, etc. For more distant genes, there is insufficient power to estimate these probabilities so AMM recommends combining these into bins. We follow the recommended binning, and then use the probabilities estimated in the original AMM paper [49, Supplementary Table 5]. AMM recommends using LDSC baseline covariates in all models, for which we used v2.3. We restricted our analysis to HapMap3 variants. The results in Figure 6**D** are the inverse-variance weighted average of the heritability enrichment estimates across our 27 genetically uncorrelated traits. These inverse-variance weighted estimates of the average enrichments do not necessarily need to average to one in contrast to the true enrichments. As such, we renormalized the estimated enrichments so that they sum to one.

### Probability of a variant being a GWAS hit for a gene correlates with *s*_**het**_

We analyzed the GWAS hits curated in our previous study [47], filtered to a set of 6,971,256 SNPs that passed quality control procedures. Importantly, this set excluded lead GWAS SNPs in LD (*r*^2^ > 0.8) with variants predicted to have protein-altering consequences, to condition on putatively non-coding trait associations. We focused on 15,591 approximately independent GWAS hits associated with our 27 uncorrelated traits, for which estimates of *s*_*het*_ for the nearest gene were available. We performed logistic regression to differentiate GWAS hits from 100,000 SNPs randomly sampled from the same 6,971,256 SNP set. The *s*_*het*_ values of the nearest genes were used as the predictor, categorized into 100 percentile bins. As in our previous work, the regression model included additional covariates: minor allele frequency (MAF), LD score, gene density, and the absolute distance to the nearest transcription start site (TSS). We also incorporated dummy variables representing 20 quantiles of each of these covariates (MAF, LD score, gene density, and distance to TSS). Results of this analysis are presented in Supplementary Figure S18. The covariate data were obtained from Mostafavi *et al*. [47].

### The number of GWAS hits near a gene correlates with trait importance

To avoid double counting GWAS hits due to LD, we restricted our analysis to approximately independent hits. For each trait, we analyzed the set of LD-clumped hits (*p* < 5 × 10^−8^, clumping *r*^2^ < 0.1) from 8,136,100 filtered SNPs provided in [47]. We then assigned each GWAS hit to the closest gene (using the midpoint of genes as released with AMM [49]). For each trait, we then correlated the number of GWAS hits assigned to each gene with our unbiased estimate of that gene’s trait importance, 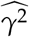 based on the LoF burden test results. To make certain that our results were not driven by differences between genes with no GWAS signal versus genes with any GWAS signal, we also computed correlations between number of GWAS hits assigned to each gene and 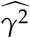 restricting to genes with at least one GWAS hit.

## Acknowledgments

We thank Terence Capellini, Emma Dann, Molly Przeworski, Daniel Richard, Catherine Tcheandjieu, Huisheng Zhu, and members of the Pritchard Lab for helpful feedback. This work was supported by the National Institutes of Health (grants R01HG011432, R01HG008140, U01HG012069 to J.K.P. and R01GM115889 to G.S.).

## Supplementary Table

**Supplementary Table 1:** List of traits and abbreviations used in the study. Table of the 209 traits used in this study with the UKB trait IDs, trait names, abbreviations used, tissue to which each trait was linked (if applicable), and an indication of whether or not the trait was included in our subset of 27 genetically uncorrelated traits (Methods).

**Figure S1:**
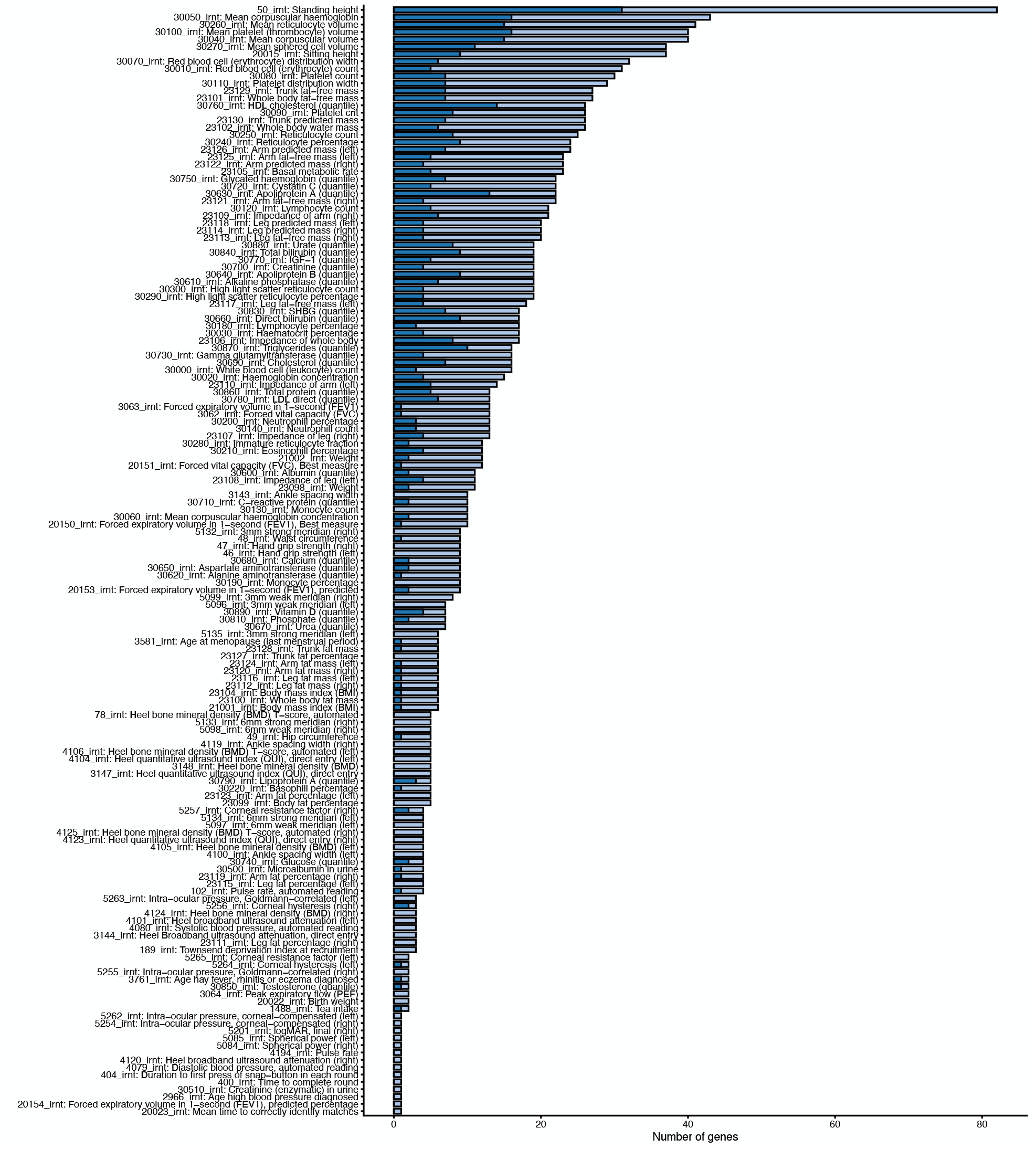
Limited GWAS overlap at top LoF burden hits. Extended version of Figure 1**C**, including all traits with at least one genome-wide significant LoF burden test. Dark blue bars correspond to genome-wide significant LoF burden test genes that also overlap a top GWAS locus (Methods). Light blue bars are genome-wide significant LoF burden test genes that do not overlap a top GWAS locus.

**Figure S2:**
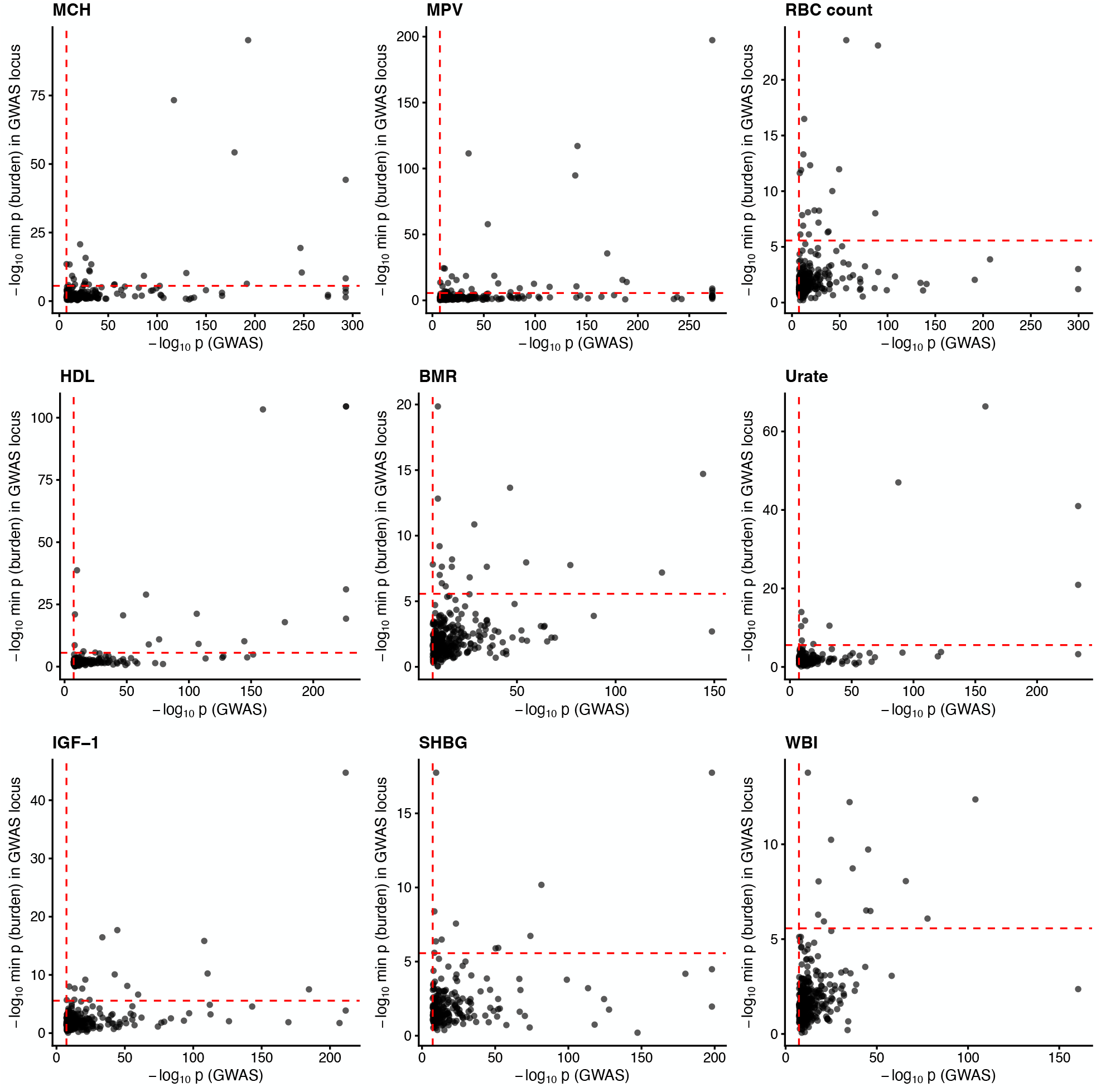
Strongest GWAS hits are not always strong LoF burden hits. Extended version of Figure 1**D**, including 9 additional traits. Each point is a significant GWAS locus (Methods). Dashed red lines are thresholds for genome-wide significance.

**Figure S3:**
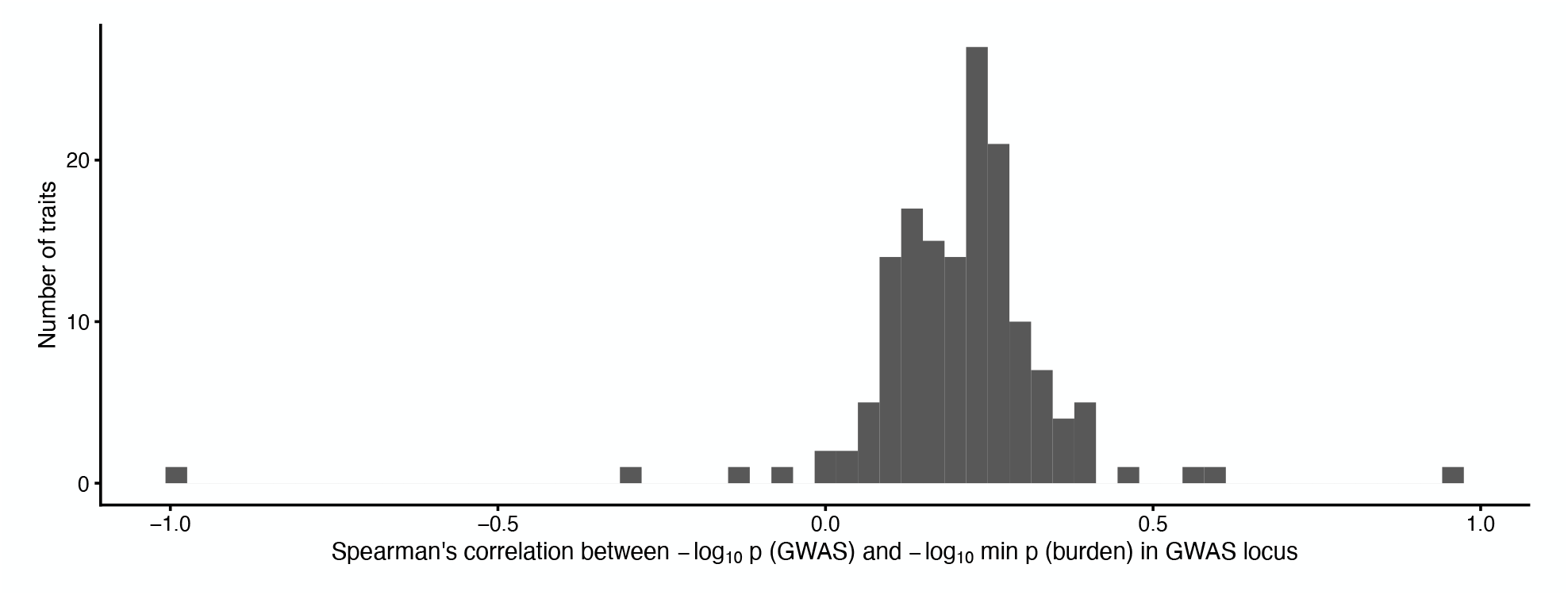
Modest correlation between GWAS and LoF burden test p-value ranks. Histogram of Spearman’s ρ between the minimum GWAS − log_10_ p-value of any variant within a given GWAS locus and the minimum LoF burden − log_10_ p-value of any gene overlapping that locus. By definition all GWAS loci contain at least one genome-wide significant variant (Methods).

**Figure S4:**
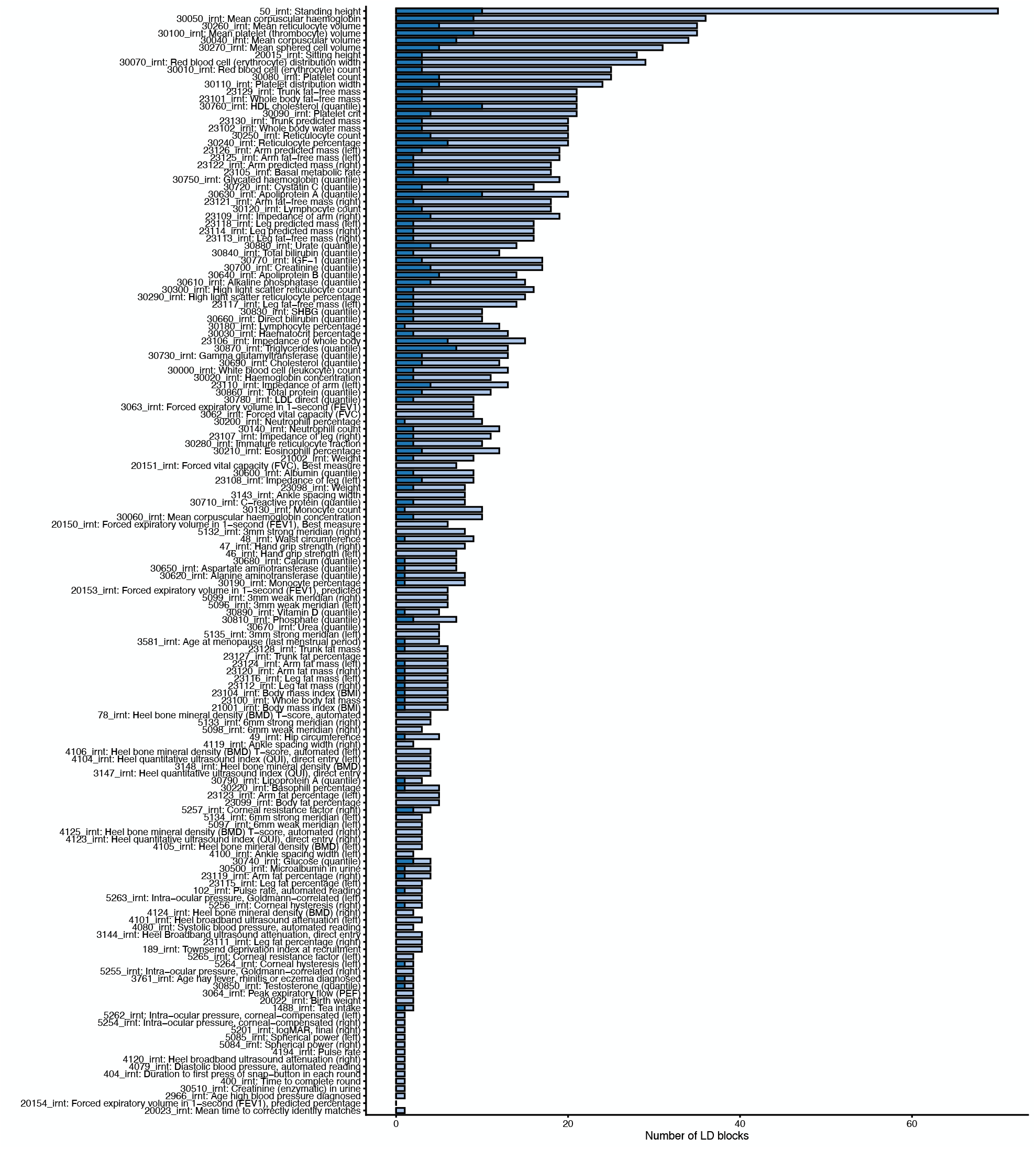
Limited GWAS overlap at top LoF burden LD blocks. Alternate version of Supplementary Figure S1 but using LD blocks instead of GWAS loci. Dark blue bars correspond to LD blocks that contain a genome-wide significant LoF burden test gene that are also top LD blocks for GWAS. Light blue bars are LD blocks containing genome-wide significant LoF burden test genes that are not also top GWAS LD blocks.

**Figure S5:**
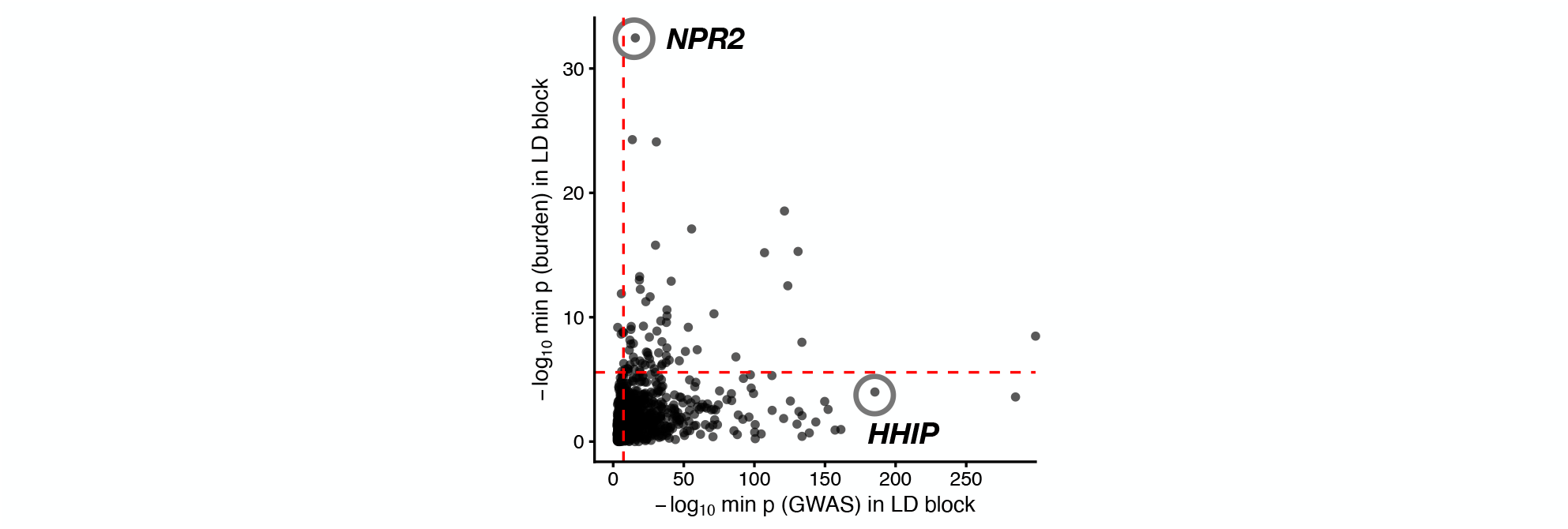
GWAS and LoF burden tests prioritize different LD blocks for height. Alternate version of Figure 1**D** but using LD blocks instead of GWAS loci. Each point is an LD block. Dashed red lines are thresholds for genome-wide significance.

**Figure S6:**
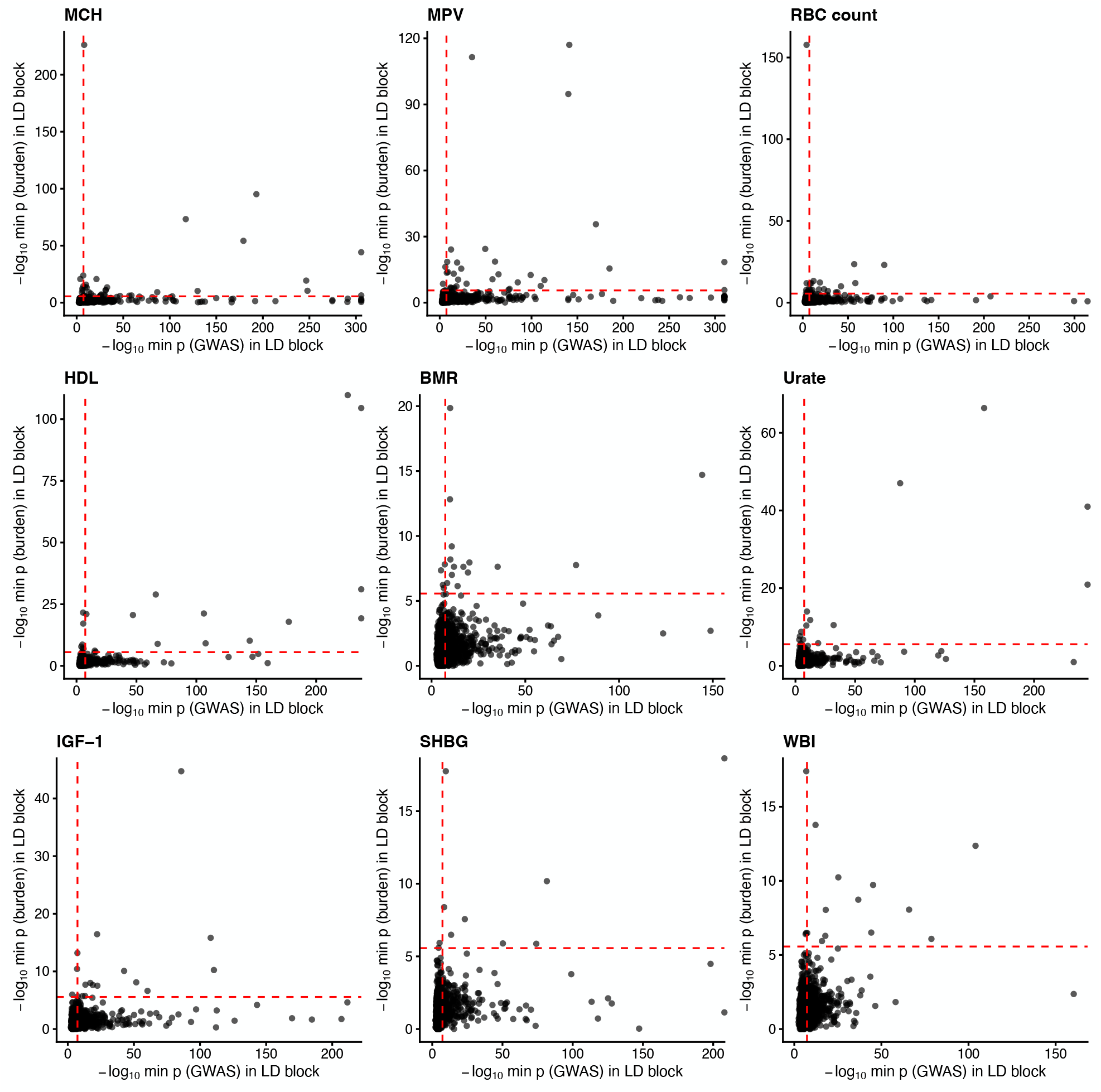
GWAS and LoF burden tests prioritize different LD blocks across traits. Alternate version of Supplementary Figure S2 but using LD blocks instead of GWAS loci. Each point is an LD block. Dashed red lines are thresholds for genome-wide significance.

**Figure S7:**
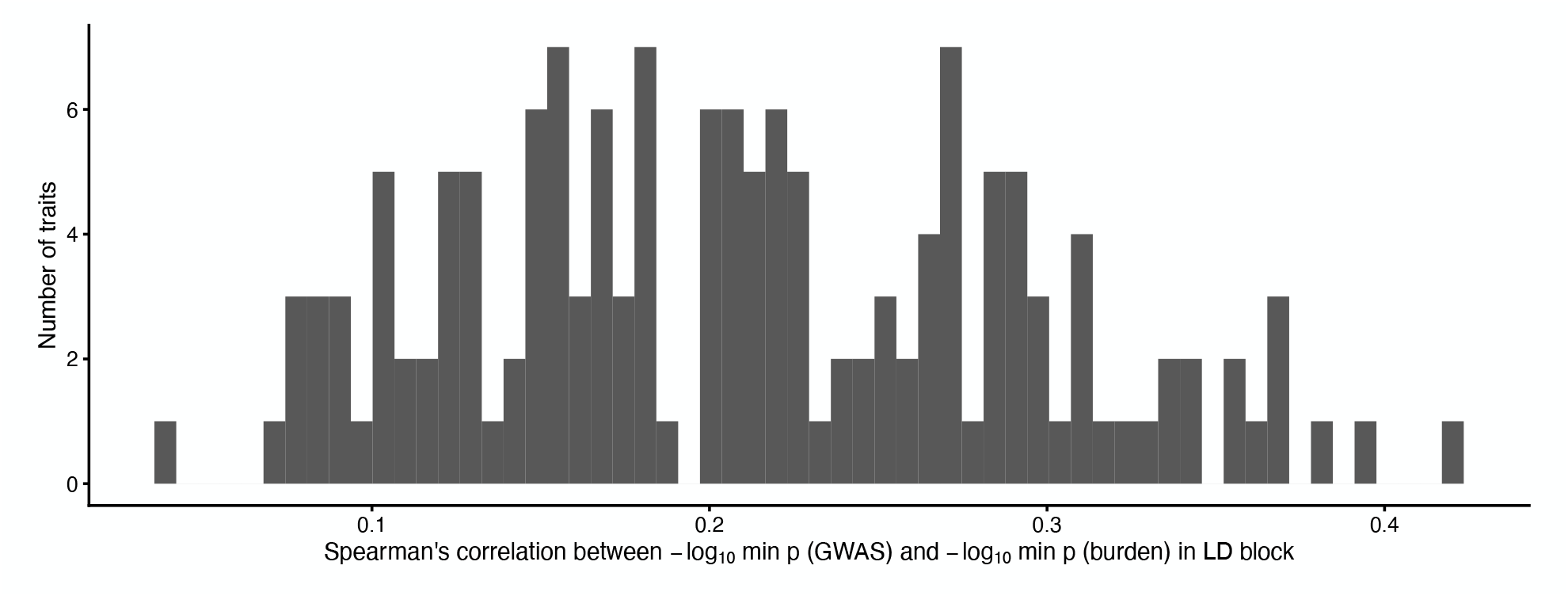
Modest correlation between GWAS and LoF burden test p-value ranks across LD blocks. Alternate version of Supplementary Figure S3 but using LD blocks instead of GWAS loci. Histogram of Spearman’s ρ between the minimum GWAS − log_10_ p-value and the minimum LoF burden − log_10_ p-value across LD blocks.

**Figure S8:**
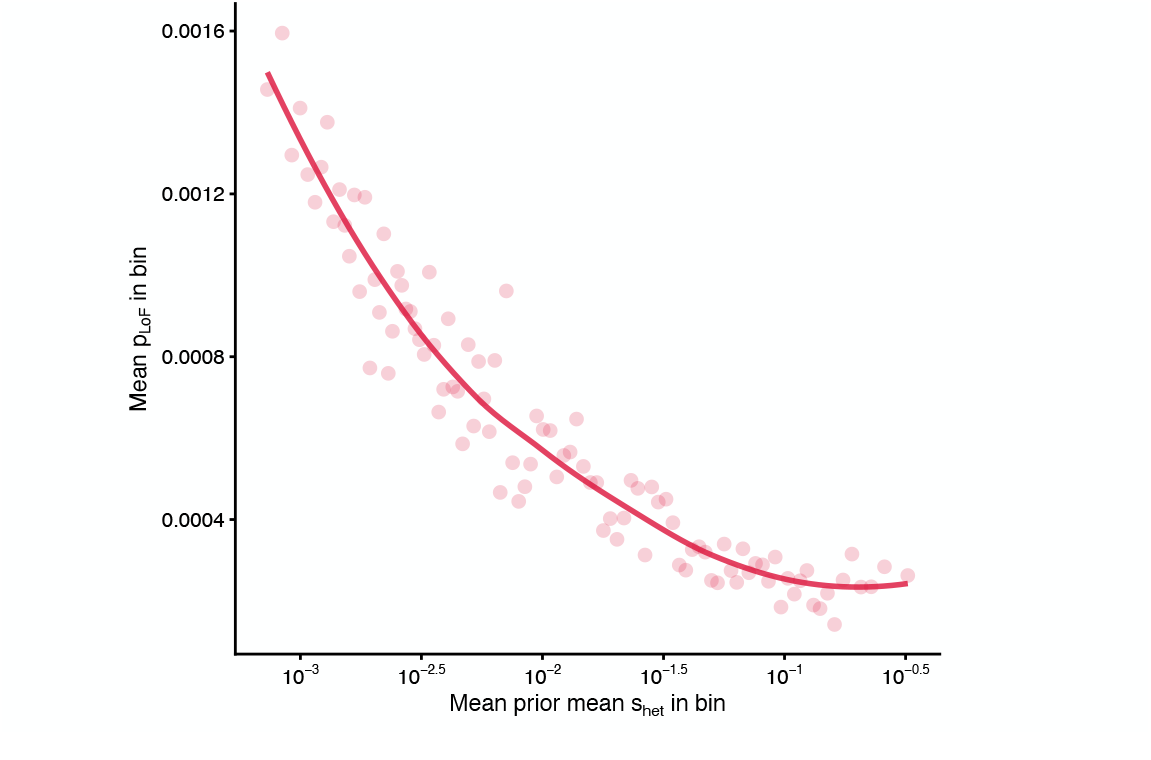
p_LoF_ and s_het_ are negatively correlated. Alternate version of Figure 3**B** but binning genes by the prior mean s_het_ as reported by [33]. These estimates are learned using GeneBayes [33], which uses frequency data across genes to learn a function mapping gene features (e.g., expression patterns across tissues) to a prior on s_het_. In the main text, we used GeneBayes posterior mean estimates, which use this learned prior for each gene along with that gene’s p_LoF_ to estimate s_het_. Here we use the prior mean, which uses the p_LoF_ data across genes to learn per-gene priors, but does not use a gene’s p_LoF_ when estimating its s_het_.

**Figure S9:**
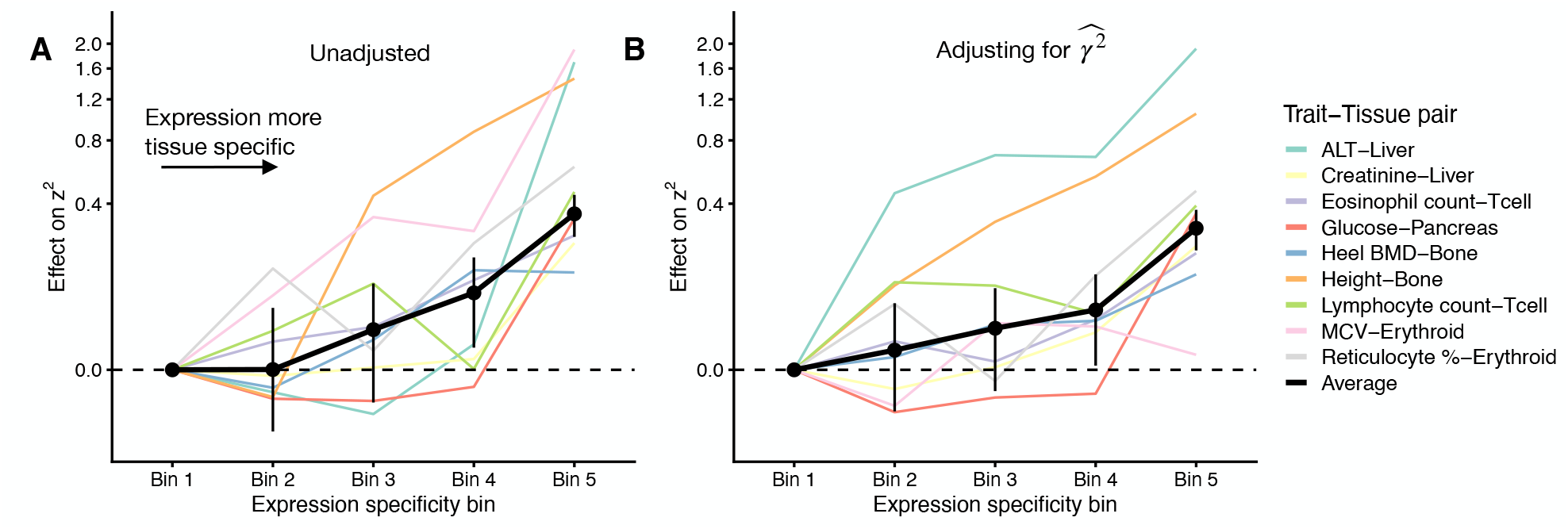
Expression specificity increases LoF burden test z-scores. For 9 trait-tissue pairs we regressed LoF burden test z^2^ for each gene on either **A**) expression specificity bin or **B**) expression specificty bin and an unbiased estimate of 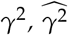. Since the 5 bins are co-linear, we report all regression coefficients relative to the effect in expression specificity bin 1. Colored lines are regression coefficients for individual trait-tissue pairs. The black line is the inverse variance-weighted average across trait-tissue pairs. The y-axes have been non-linearly transformed.

**Figure S10:**
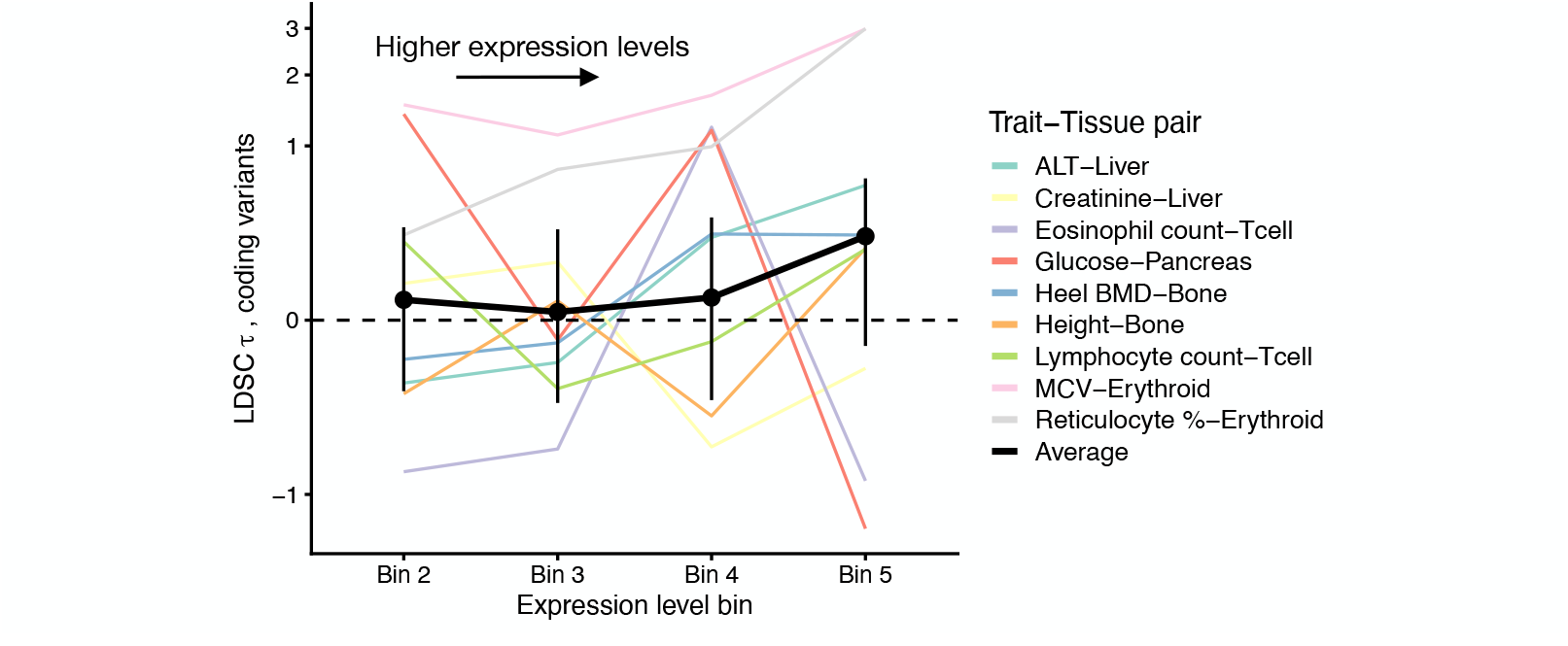
Expression levels do not play a large role in GWAS coding heritability. S-LDSC analysis results for 9 trait-tissue pairs. Result are reported in terms of τ, a measure of heritability enrichment. Variants are binned by the expression level (as measured by TPM) of the corresponding gene. Since the 5 bins are co-linear, we drop the bin 1 annotation and only report results for the remaining bins. These results are from a joint analysis including both expression specificity and expression level bins as covariates. Colored lines are τ estimates for individual trait-tissue pairs. The black line is the inverse variance-weighted average across trait-tissue pairs. The y-axis has been non-linearly transformed.

**Figure S11:**
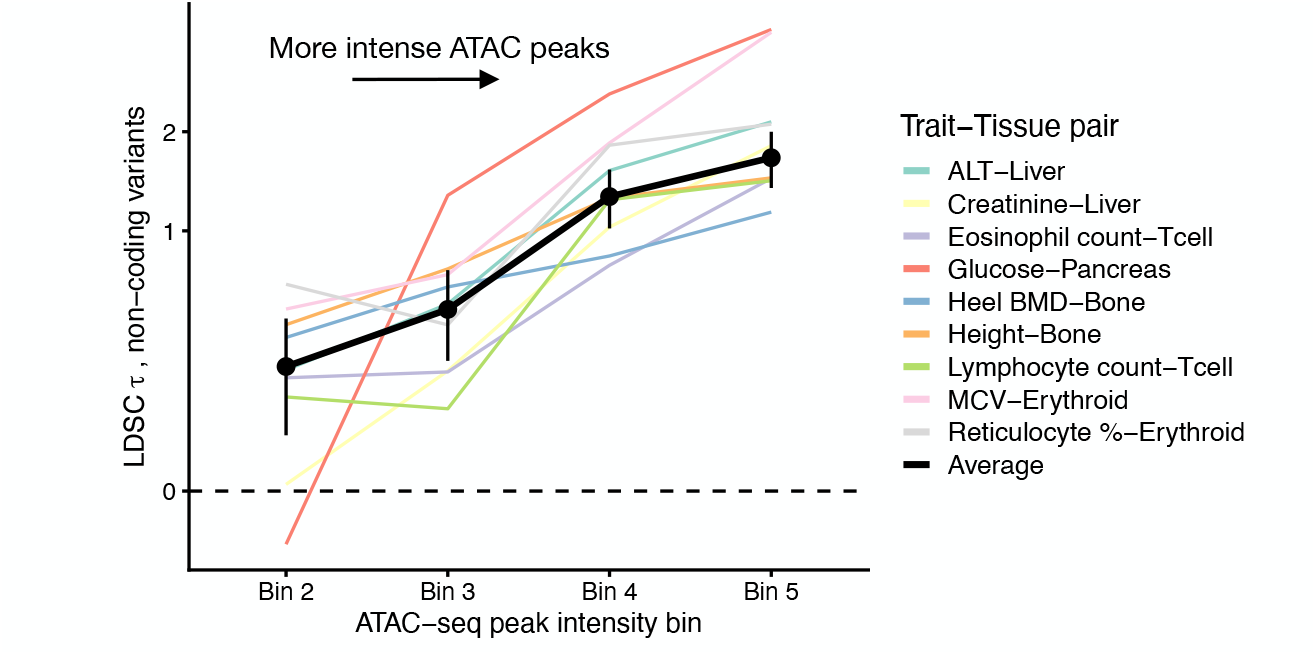
ATAC peak intensity increases heritability explained. S-LDSC analysis results for 9 trait-tissue pairs. Result are reported in terms of τ, a measure of heritability enrichment. Variants are binned by the intensity of their ATAC-seq peaks (Methods). Since the 5 bins are co-linear, we drop the bin 1 annotation and only report results for the remaining bins. These results are from a joint analysis including both ATAC specificity and ATAC intensity bins as covariates. Colored lines are the τ estimates for individual trait-tissue pairs. The black line is the inverse variance-weighted average across trait-tissue pairs. Note that the y-axis has been non-linearly transformed.

**Figure S12:**
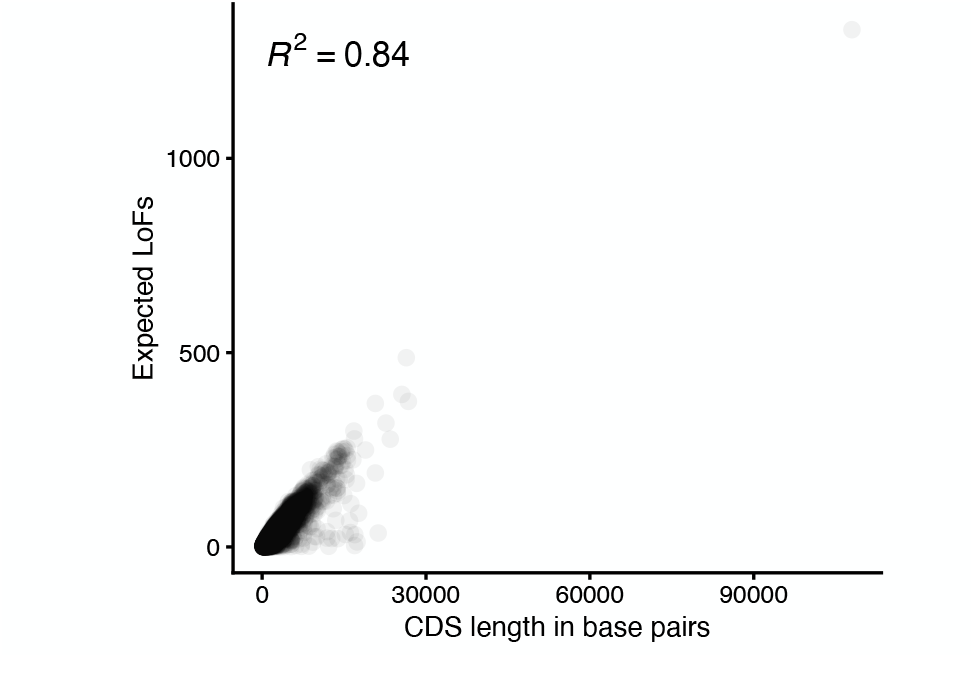
CDS length and expected number of unique LoFs are highly correlated. Scatter plot of the expected length in base pairs for the canonical CDS for each gene (Methods) and the expected number of unique LoFs as computed by gnomAD [43]. The overall correlation is high (Pearson’s r^2^ = 0.84).

**Figure S13:**
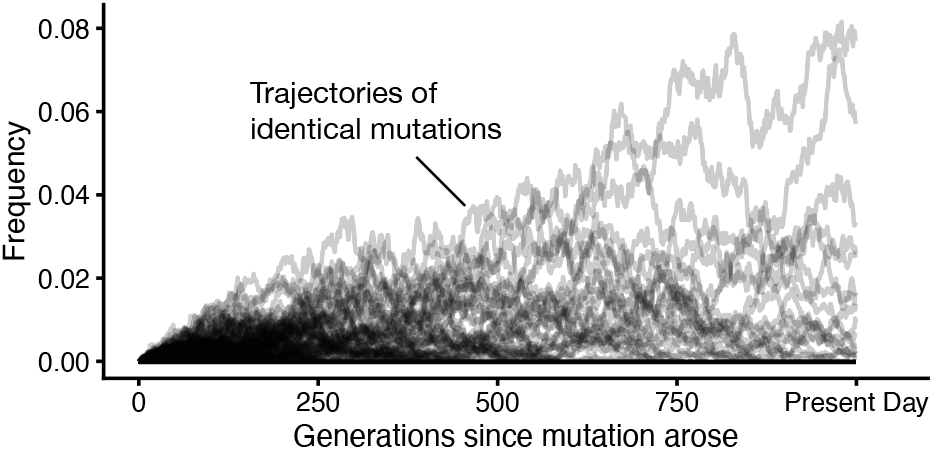
Genetic drift generates variance in allele frequencies. 10,000 frequency trajectories of identical mutations simulated under the Discrete-Time Wright-Fisher model. Trajectories were simulated assuming no mutation, an s_het_ of 10^−3^, no fitness consequences in homozygotes, and a population size of N_e_ = 10,000. All mutations were assumed to arise 1,000 generations before present.

**Figure S14:**
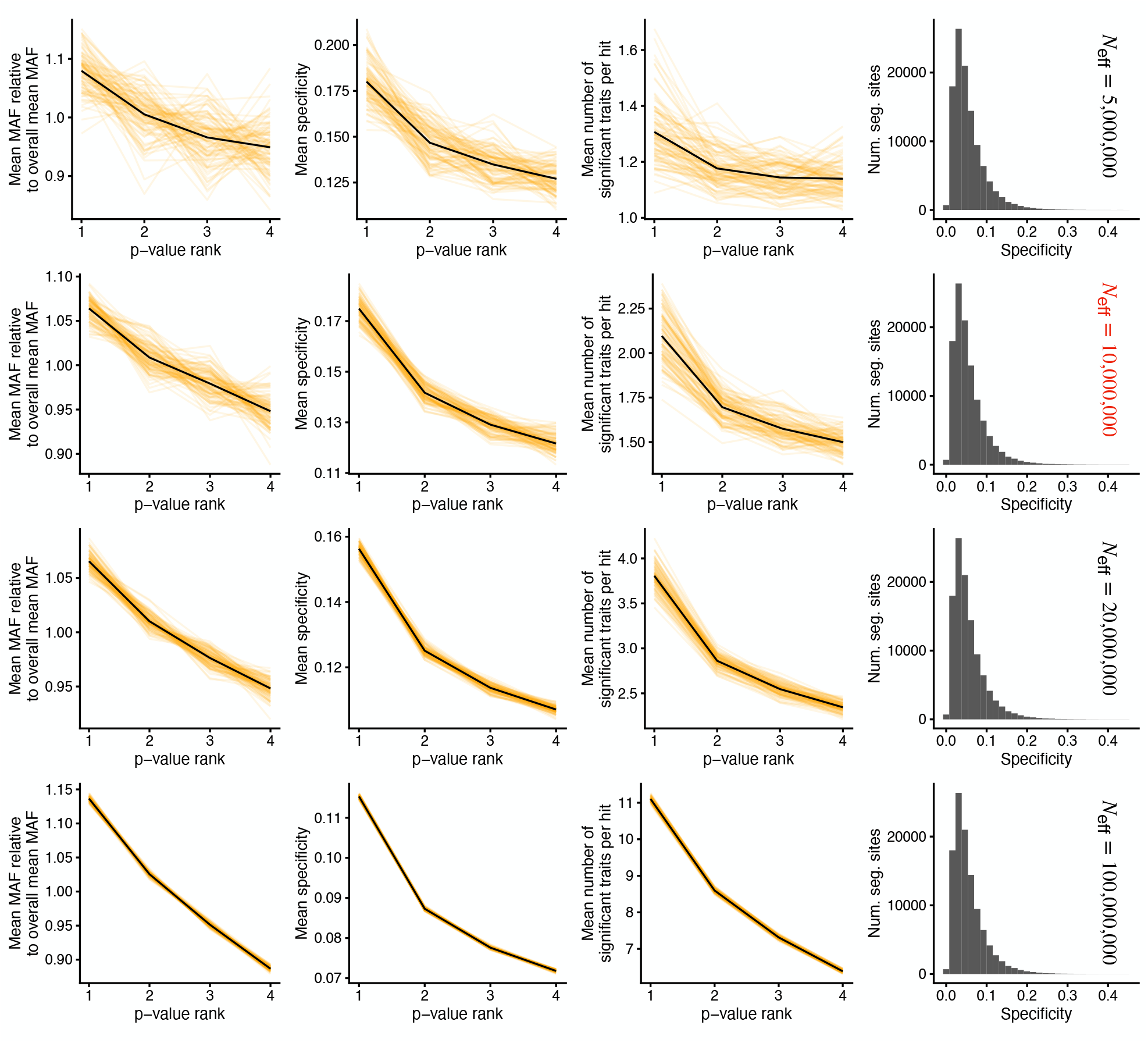
Robustness of apparent pleiotropy to simulation parameter N_**eff**_. Analogous to Figure 5**F**-**H**, but with varying N_eff_ (see Methods for definition), while holding all other simulation parameters fixed to the values used in the main text. Results from individual population genetic simulations are in orange, and the mean across simulations is in black. The histograms show the distribution of trait specificity, Ψ_V_, across segregating sites for a single simulation. N_eff_ does not affect the distribution of effect sizes, and so these are the same across values of N_eff_. The value of N_eff_ used in the main text is in red.

**Figure S15:**
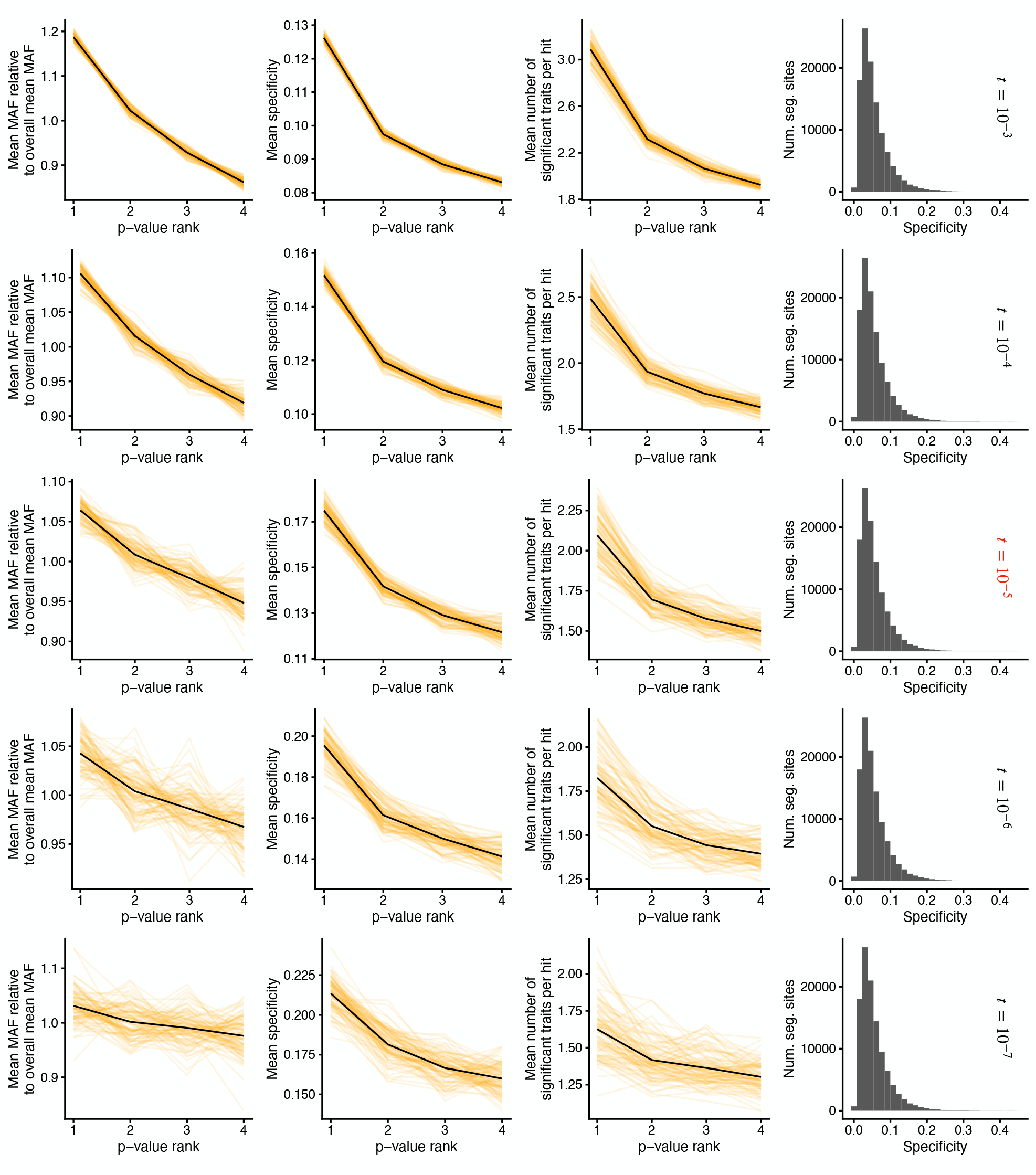
Robustness of apparent pleiotropy to simulation parameter t. Analogous to Figure 5**F**-**H**, but with varying t (see Methods for definition), while holding all other simulation parameters fixed to the values used in the main text. Results from individual population genetic simulations are in orange, and the mean across simulations is in black. The histograms show the distribution of trait specificity, Ψ_V_, across segregating sites for a single simulation. t does not affect the distribution of effect sizes, and so these are the same across values of t. The value of t used in the main text is in red.

**Figure S16:**
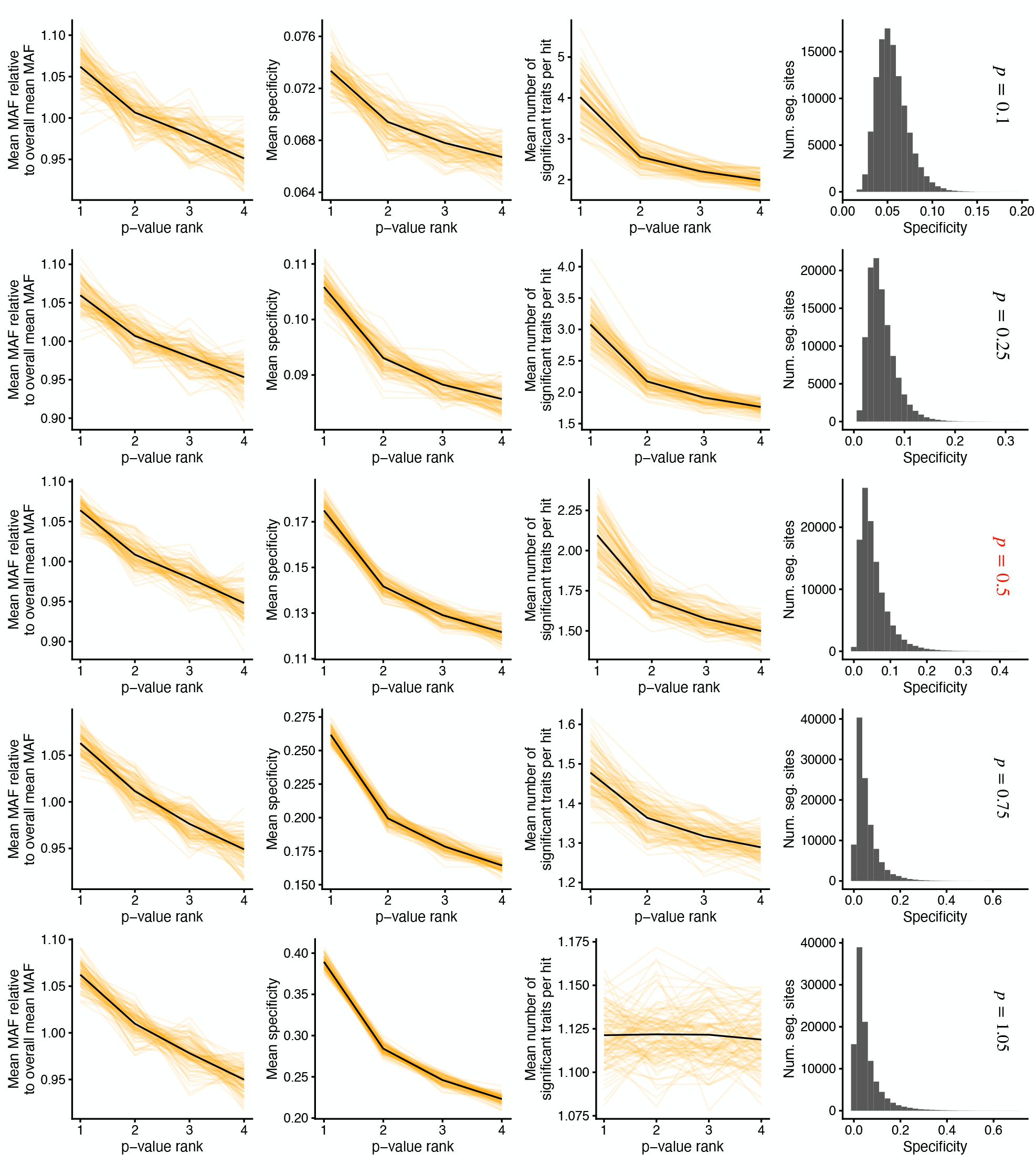
Robustness of apparent pleiotropy to simulation parameter p. Analogous to Figure 5**F**-**H**, but with varying p (see Methods for definition), while holding all other simulation parameters fixed to the values used in the main text. Results from individual population genetic simulations are in orange, and the mean across simulations is in black. The histograms show the distribution of trait specificity, Ψ_V_, across segregating sites for a single simulation. The value of p used in the main text is in red.

**Figure S17:**
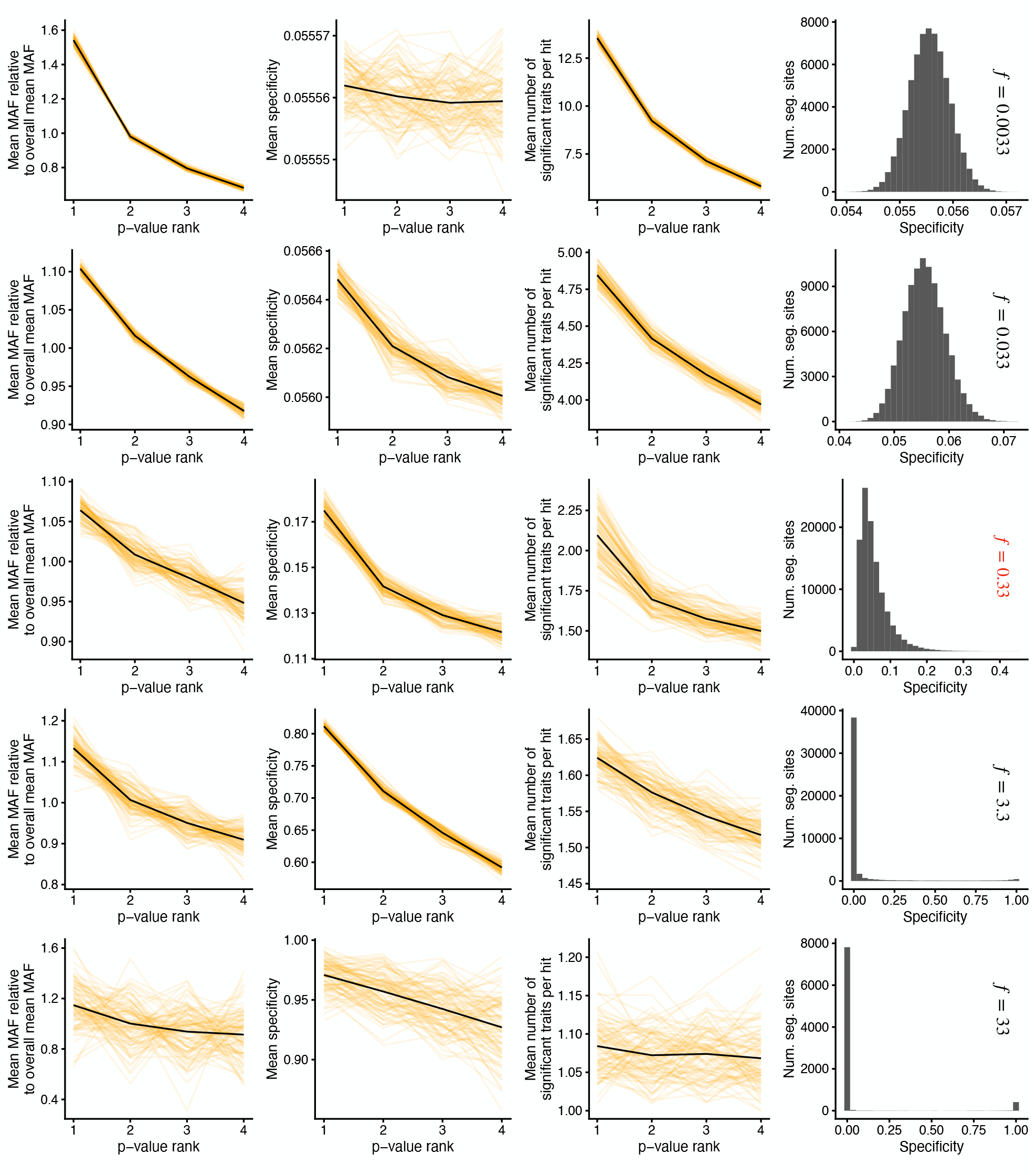
Robustness of apparent pleiotropy to simulation parameter f . Analogous to Figure 5**F**-**H**, but with varying f (see Methods for definition), while holding all other simulation parameters fixed to the values used in the main text. Results from individual population genetic simulations are in orange, and the mean across simulations is in black. The histograms show the distribution of trait specificity, Ψ_V_, across segregating sites for a single simulation. The value of f used in the main text is in red.

**Figure S18:**
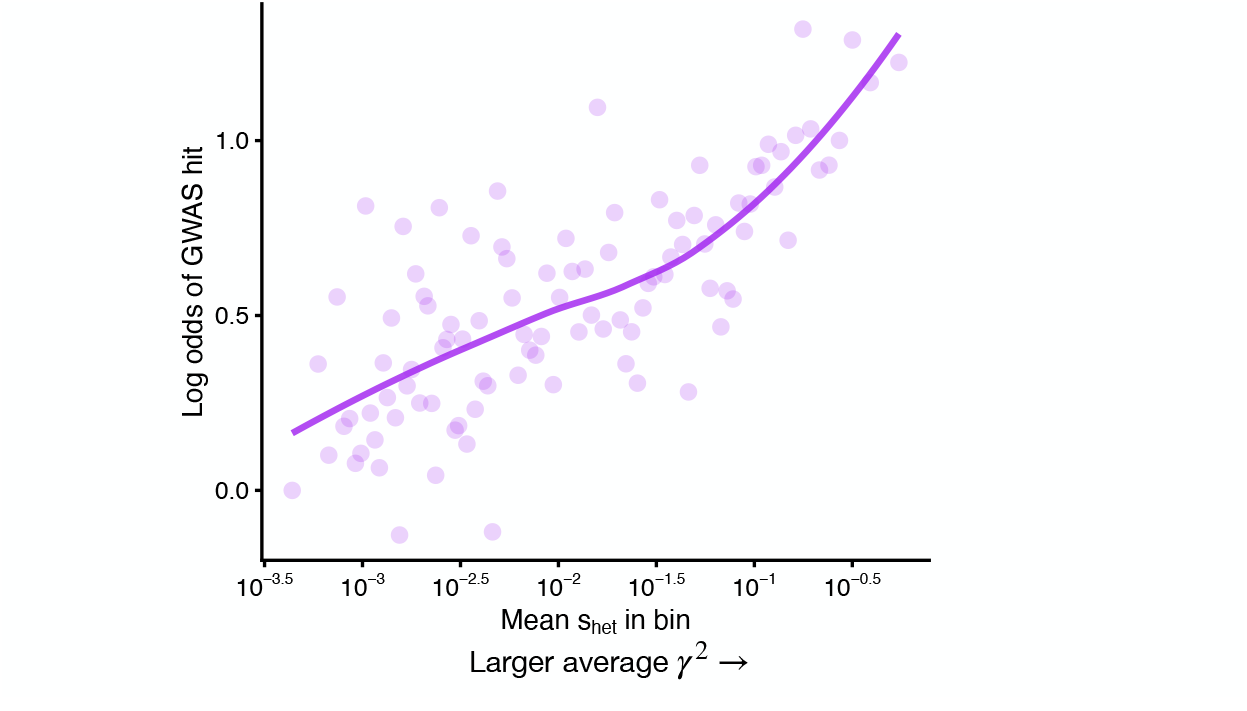
Probability of a variant being a GWAS hit is correlated with s_**het**_. Logistic regression coefficients for s_het_ percentile categories in a model that predicts whether a variant is a GWAS hit or not including various covariates such as distance to transcription start site (Methods). Each bin contains approximately 184 genes. The trend line is fit using LOESS.

**Figure S19:**
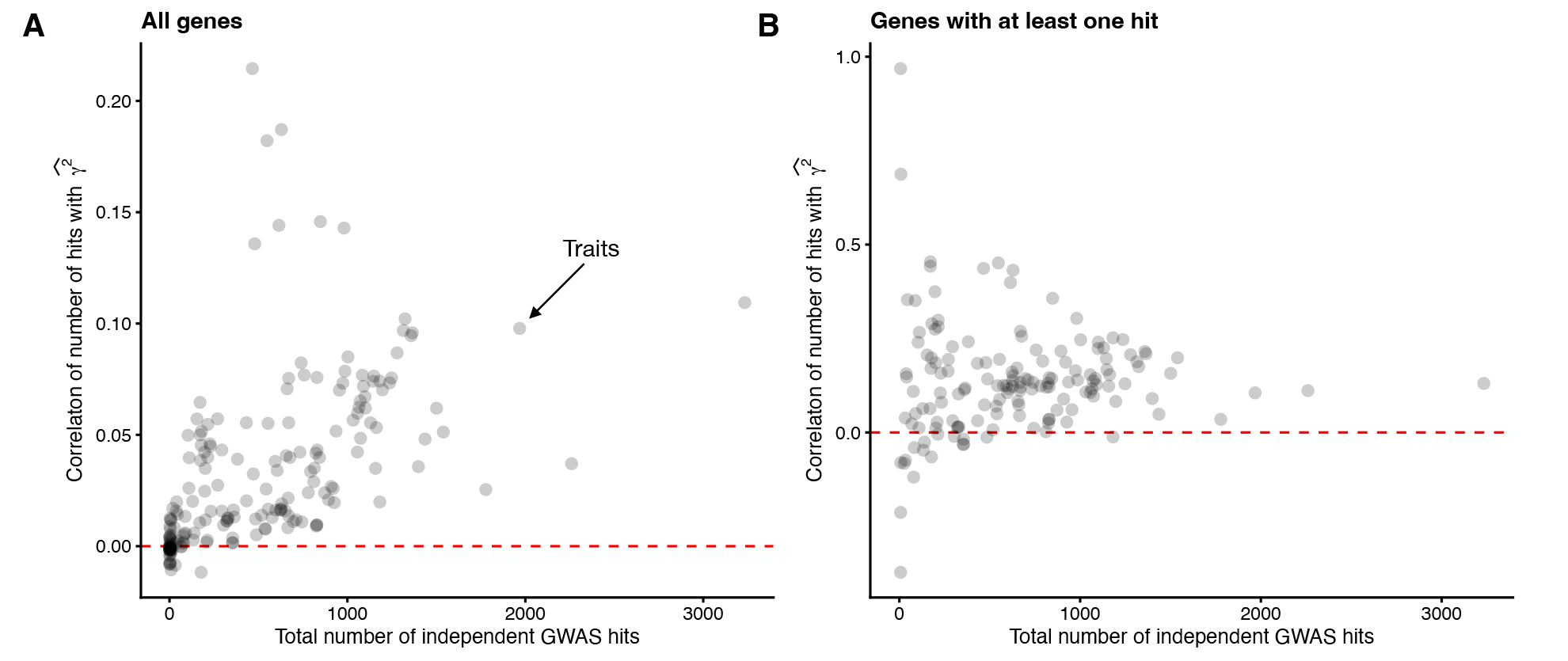
Number of GWAS hits is predictive of γ^2^. Scatter plots of the correlation across genes between the number of independent GWAS hits and an unbiased estimate of 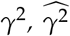, against the total number of independent GWAS hits. **A**) Correlation across all genes. Traits with more independent hits tend to have a higher correlation between number of hits and 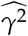. **B**) To make sure that the correlations in panel **A** were not driven just by presence or absence of any GWAS hits, we computed correlations between number of GWAS hits and 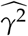 for only those genes with at least one GWAS hit. In both panels, it should be noted that 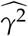 is generally a very noisy estimate of γ^2^. This will drive the plotted correlations to be much lower than the true correlation between the number of GWAS hits and the unobserved true values of γ^2^.

## A A mathematical model of association studies

In this appendix we describe our mathematical model of association studies and clarify the relationship between population, quantitative, and statistical genetics concepts. Our results rely primarily on the work of [65] and [38].

### Main theoretical results

We begin with an additive model of phenotypes:

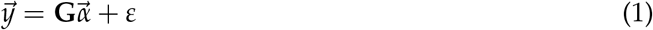

where 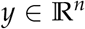 is the vector of phenotypes of the *n* individuals included in the study, 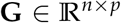 is a matrix containing the centered, but not scaled genotypes of the *n* individuals at the *p* causal loci,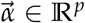 is the vector of the effects of the “1” alleles at each causal locus on the phenotype (usually denoted by *β* in the literature, but we reserve *β* for the effect of variants on genes), and 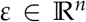 is a random vector representing unobserved noise. This additive model is rather simple, but is generally well-supported as a good approximation for many complex traits [35] and is a common assumption across statistical genetics [76–81]. We assume that the causal variants are unlinked in the population, and that *p* is large enough such that the amount of heritability contributed by any single site is ≪ 1. We also assume that the phenotype is measured in units so that its variance across individuals is 1.

Before proceeding further, we note that there are several distinct sources of randomness in Equation 1. First, the environmental noise affecting each individual is random, which is made explicit by *ε* being a random variable. Second, the choice of which individuals are included in the GWAS is random: one could imagine sampling a separate cohort from the same population and obtaining individuals with different genotypes and phenotypes. Third, the genotypes and phenotypes in the population itself are the result of the fundamentally random process of evolution. Note that this is distinct from just obtaining another GWAS cohort. Taking two large random samples from the same population will result in the two cohorts having almost perfectly correlated allele frequencies, but replaying the tape of evolution would result in alleles having completely different frequencies. Throughout, we will try to be clear about which sources of randomness we are averaging over or point out when a particular source of randomness is negligible.

Given our assumptions, it has been shown [65] that conditioned on the GWAS sample, 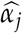, the estimate of the *j*^th^ marginal effect,*α*_*j*_, is asymptotically Normally distributed:

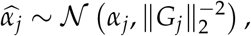

where *G*_*j*_ is the *j*^th^ column of **G** (i.e., the vector of genotypes at the *j*^th^ causal variant). Similarly, the standard error of 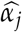 estimated by GWAS is 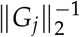. Therefore, the z-score, for variant *j*, 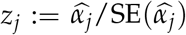, reported by GWAS is also asymptotically Normally distributed:

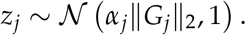

Equivalently, we can write *z*_*j*_ as the sum of a constant and a standard Normal random variable, say *u*_*j*_:

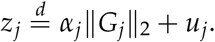

We therefore have that the GWAS test statistic, *z*^2^ (sometimes referred to as the chi-squared statistic as it is asymptotically chi-squared distributed under a null hypothesis of no effect of the variant on the trait), is

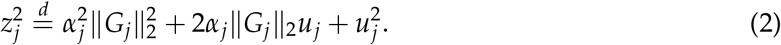

Note that *u*_*j*_ is determined by the environment randomness in *ε*. Averaging over the randomness in *ε*, which we denote by 𝔼_*ε*_ we obtain that the expected 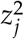 statistic is

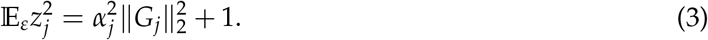

Note that the right-hand side of equation 3 depends on the genotypes of the individuals included in the GWAS. To relate this to population quantities, we assume that the individuals included were sampled randomly from the population (and hence independently from the phenotype), and that the population is at Hardy-Weinberg equilibrium. In such a case, the variance of a randomly sampled genotype with population frequency *f*_*j*_ is 2 *f*_*j*_(1 − *f*_*j*_), which can be seen by considering the genotype as the sum of two randomly chosen haploids, which are each independent Bernoulli(*f*_*j*_) random variables. Then, we see that averaging over this sampling process, which we denote by 𝔼_**G**_, results in

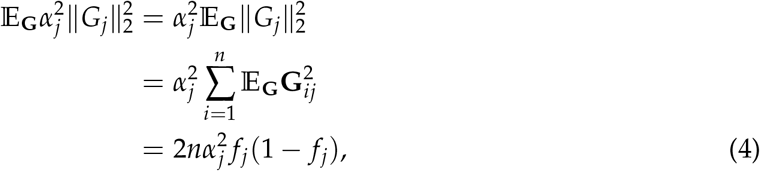

where the first line follows from our assumption that individuals were chosen independently of the phenotype, and the final follows from the fact that we are considering centered genotypes, so 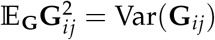.

Plugging equation 4 into equation 3 we obtain that averaging over environmental randomness and the individuals included in the GWAS, we obtain

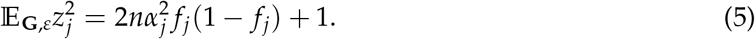

As a result, under our assumptions the average strength of association for the *j*^th^ variant in GWAS is determined by 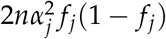.

By a similar argument, as *n* → ∞,

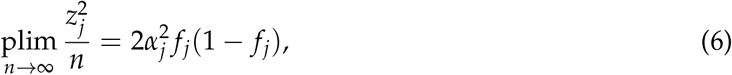

and so an infinitely powered GWAS would prioritize variants exactly by 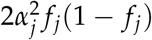. To make this argument rigorous, note that in equation 2, **G**_ij_ is bounded so ∥*G_j_*∥2 is 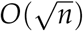 and *u_j_* is *O*_*p*_ (1),so only the 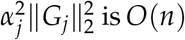. Then, again because 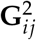 is bounded, 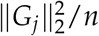 converges in probability to its mean, 2 *f*_*j*_(1 − *f*_*j*_) by the weak law of large numbers.

From the perspective of population genetics, the frequencies appearing equations 5 and 6 are the result of evolution, which is itself a random process. As such, we might consider how such sites typically behave under an evolutionary model. In this section, we assume that selection is sufficiently strong against new mutations that we are in the mutation-selection balance regime. See Appendix C for the general case. For the subsequent discussion, we will assume without loss of generality that the “1” allele at each locus is the minor allele.

In this regime, selection is strong enough that the minor allele will be at a low enough frequency such that 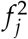 is essentially negligible, so *f*_*j*_(1 − *f*_*j*_) ≈ *f*_*j*_. Then, 𝔼 *f*_*j*_ = *µ*/*s*_het_, where *µ* is the mutation rate, and *s*_het_ is the strength of selection against heterozygotes [32, equation (3.9)]. Intuitively, alleles enter the population at a rate of 2*Nµ*, where *N* is the population size, and are removed from the population at a rate of *s*_het_ in the roughly 2*N f*_*j*_ heterozygous individuals. At equilibrium, these forces must cancel, resulting in *f*_*j*_ ≈ *µ*/*s*_het_. Plugging this result into equation 5, we obtain

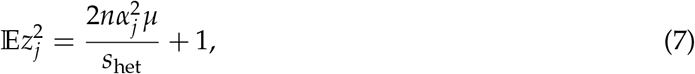

where now the expectation is taken over the environmental randomness, the randomness in the composition of the GWAS cohort, and the evolutionary process.

Finally, selection must be acting upon a variant due to its effects on some phenotypes. In Appendix D we show that under a model of stabilizing selection on *T* traits measured in appropriate units, 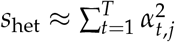, where *α*_*t*,*j*_ is the effect of the *j*^th^ variant on the *t*^th^ fitness-relevant trait. Substituting this result in equation 7, we obtain our main result

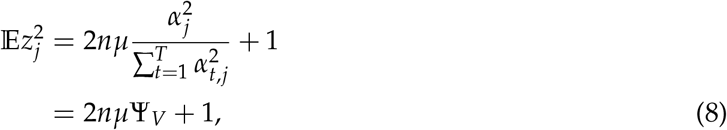

where Ψ_*V*_ is the trait specificity of the variant as defined in the main text.

### LoF burden tests

To extend these results to LoF burden tests, the only part that needs modification is our analysis of 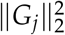, which in burden tests is a “burden genotype” instead of a standard genotype. Suppose there are *L* potential LoF positions in a gene. An individual’s burden genotype for that gene is then 1 if they have an LoF allele at any of those *L* positions. Let *f*_*𝓁*_ be the population frequency of the LoF allele at position 𝓁 within the gene. Considering a haplotype chosen randomly from the population, the probability that it does not contain an LoF allele is

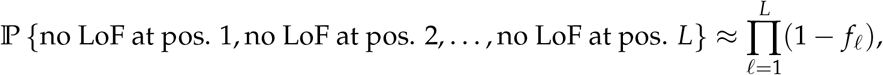

where the approximation is reasonable because LoFs tend to be extremely rare [43] and rare variants tend to be essentially independent [82]. We then continue this line of approximation, by noting that if LoFs are rare then any quadratic or higher order terms in the LoF frequencies at one or more positions are negligible. As a result,

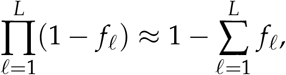

and 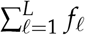 can be interpreted as the aggregate frequency of LoF variants. Assuming that this aggregate frequency is also small, we can then assume that it is unlikely to observe an individual with an LoF allele on both of their haplotypes. We can therefore approximate individual *i*’s burden genotype for gene *j*, **G**_*ij*_, as the sum of two Bernoulli 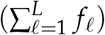 random variables. The variance of this sum is then

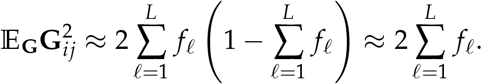

Using this result in place of equation 4, and noting that we write *γ*_*j*_ for *α*_*j*_ in the case of LoF burden tests, results in

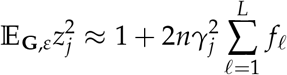

and

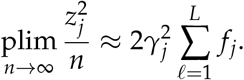

Finally, if we assume each of these *L* positions are under mutation selection balance with mutation rates *µ*, then

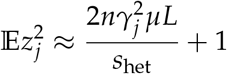

giving our main result for burden tests,

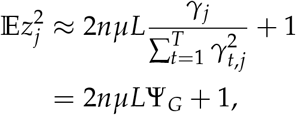

where Ψ_*G*_ is the trait specificity of the gene as defined in the main text.

### Relationship between *z*^2^, *h*^2^, and − log *p*-value

We end this section by noting that 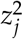 is closely related to both the heritability explained by that variant, 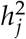, as well as the the −log *p*-value returned by an association test. We will focus on GWAS in this subsection for clarity, but the results also apply to LoF burden tests.

First, note that 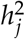 is defined (under our assumption of independent causal variants) as 2*α*^2^ *f*_*j*_(1 − *f*_*j*_) [32, equation (6.20)]. This is exactly the expected value (averaging over the environmental noise and GWAS cohort composition) of 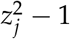 as seen in equation 5. That is,

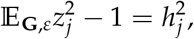

and so ranking variants based on 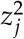 is, in expectation, equivalent to ranking based on contribution to heritability.

Next, note that under a null hypothesis where *z* has a standard Normal distribution (i.e., the null used in GWAS),

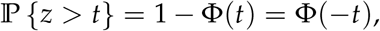

where Φ(*t*) is the standard Normal cumulative distribution function. This implies that for *t* ≥ 0,

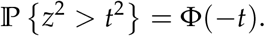

This means that the reported *p*-value is related to *z*^2^ by

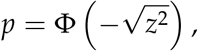

which implies that

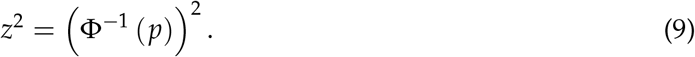

For small *p*, Φ^−1^ is asymptotically [83]

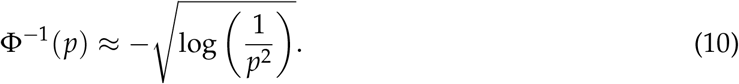

Using the approximation in equation 10 in equation 9 we obtain that for small *p*-values, *p* is related to *z*^2^ by

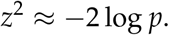

Therefore for large values of *z*^2^, it is essentially the same as twice the natural log *p*-value. In GWAS it is standard to visualize and discuss − log_10_ *p*-values, but these are just a constant scaling times the natural log *p*-values:

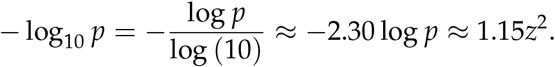

Because of these results, we freely switch between discussing average *z*^2^ statistics, contributions to heritability, and − log *p*-values in the main text.

## B model of context specificity to explain variant specificity

The main goal of this appendix is to build a model of how variants affect phenotypes that incorporates both the context dependence of variants and gene-level pleiotropy. Specifically, we want to provide an analytical expression for *variant specificity*, Ψ_*V*_, in terms of *gene specificity*, Ψ_*G*_, and a term that can be interpreted as “context specificity” under such a model.

Throughout, we will consider a biologically-inspired model of how a variant affects a trait. We assume that a variant affects traits by changing the “activity” of a gene in one or more “contexts”, and this gene affects traits by its activity in these “contexts”. We consider the “activity” of a gene as the ability of the gene to perform its physiological task in a given context. Similarly, the notion of “context” is completely abstract, but one could think of a context as being a specific cell type or tissue or perhaps an even more specific condition, such as a certain cell type at certain point of development when exposed to a certain stimulus. As such, we would say that a missense variant that totally disrupts protein folding would have a large negative effect on activity across all contexts, while a regulatory variant in a tissue-specific regulatory region may have a positive or negative effect on activity by respectively increasing or decreasing the expression of the gene in that context. Finally, we consider that traits have some impact on fitness, and thus affect the frequency of the variant dependent on that variant’s effects on the gene in different contexts and the effects of the gene on different traits in each context.

Our full model is summarized in Figure S20. We assume some number of contexts, *C*, and a number of traits *T*. We denote the effect of a variant on the activity of the gene in context *c* as *β*_*c*_, and the effect of the gene in context *c* on trait *t* as *γ*_*ct*_. That is, *β* is a property of the variant and *γ* is a property of the gene. We then define the effect of the variant on trait *t* as 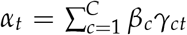. In Appendix D, we show that under mild assumptions the strength of selection in heterozygotes is equal to the sum of squared effects across all traits,

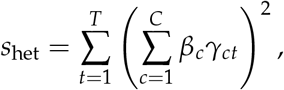

and hence under mutation selection balance, the relevant quantity for the average strength of association under this model is the trait specificity of the variant,

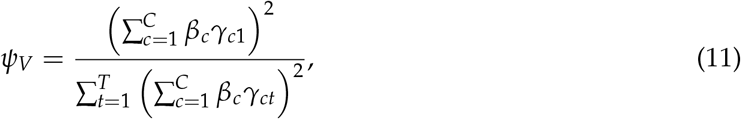

as we argued in Appendix A.

Analogously, we can consider the trait specificity of the gene, Ψ_*G*_, which under our model is

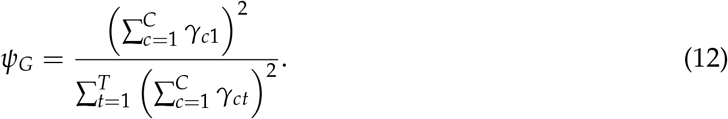

The crux of this document is defining a *context specificity factor, F*_*C*_, that captures how much more specific a variant is than the gene through which it acts. We can implicitly define *F*_*C*_ by

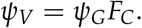

**Figure S20:**
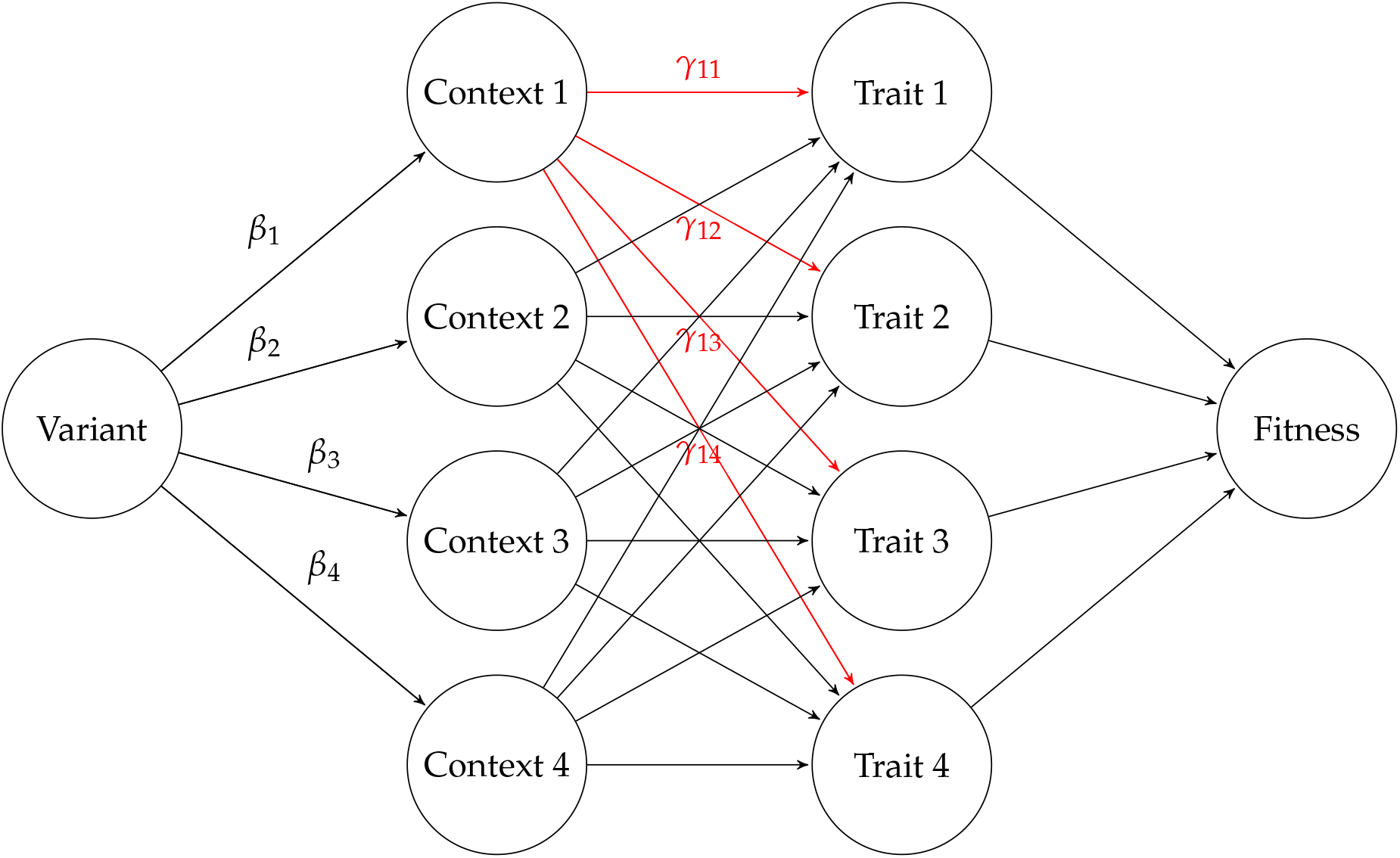
The most general model we will consider. A variant has effects in different contexts, and a gene determines how each context affects each trait.

That is,

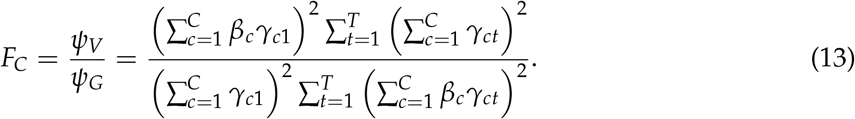

In this fully general model, the expression for *F*_*C*_ is difficult to interpret. As such, we seek a simpler model to gain some intuition. In contrast to the full model, we will now assume a simpler model where contexts and traits have a one-to-one mapping. That is, each trait is affected exclusively by a single context. We further assume that the gene either has no effect on a given trait, or has an effect of *γ*, independent of the trait. Similarly, we assume that a variant either has an effect on the activity of a gene in a context of *β* independent of the context, or has no effect on that context. This model is summarized in Figure S21.

Under our toy model the equations for *ψ*_*V*_, *ψ*_*G*_, and *F*_*C*_ simplify considerably. Let C_*V*_ be the set of contexts for which the variant has non-zero effects on the gene, and let _*G*_ be the set of contexts in which the gene has non-zero effects on the corresponding traits. Assuming that the variant affects the focal context and the gene has an effect in the focal context on the focal trait, we have

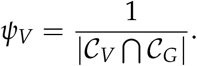

In words, *ψ*_*V*_ is 1 over the number of contexts where both the variant and gene have an effect. Similarly,

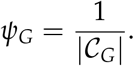

**Figure S21:**
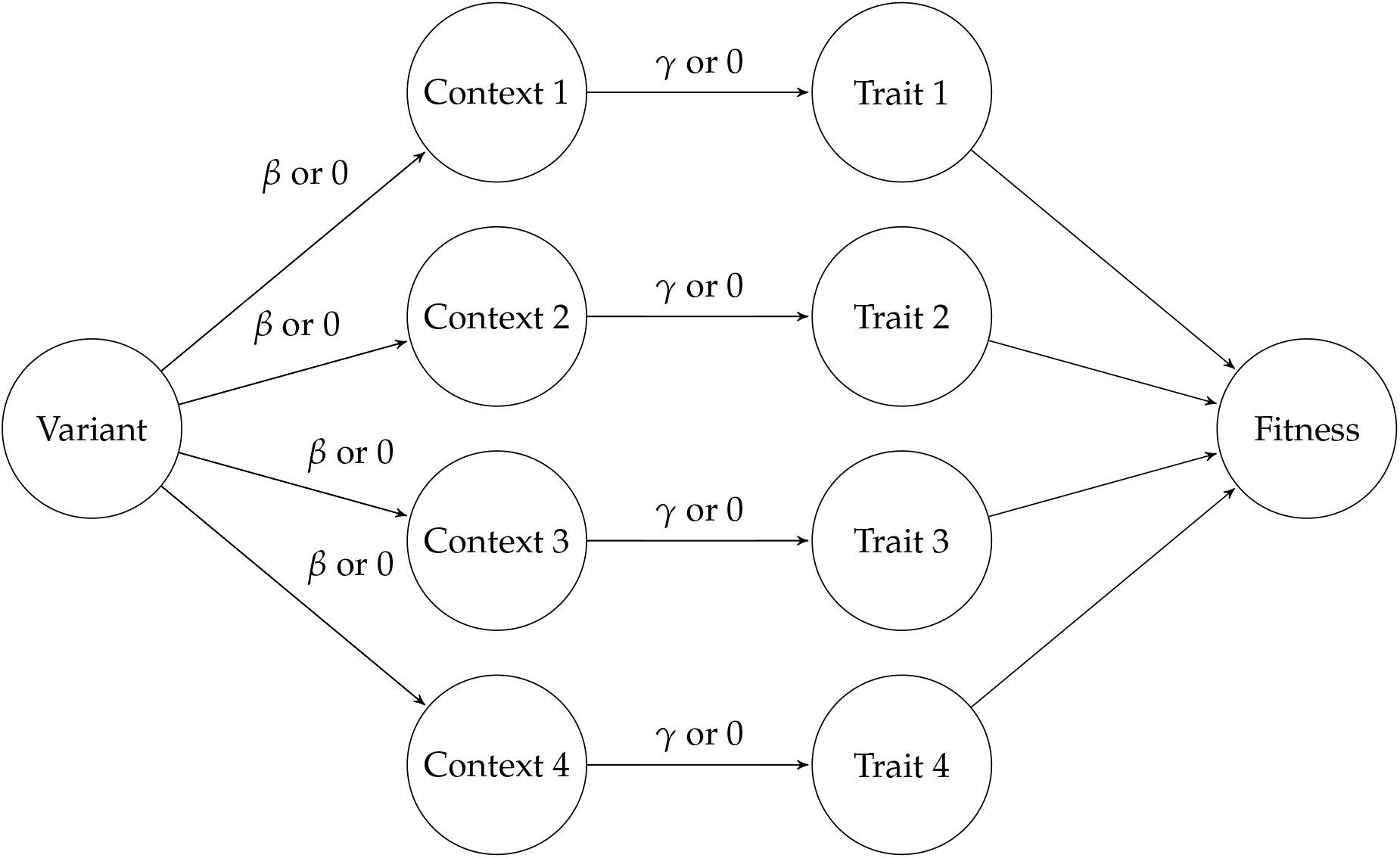
The simplest model we will consider. There is a one-to-one correspondence between contexts and traits. Variants either do or do not affect each context, and for a given gene each context may or may not affect its corresponding trait.

Again, *ψ*_*G*_ is 1 over the number of contexts where the gene has an effect. Equivalently, *ψ*_*G*_ is 1 over the number of traits that the gene affects.

Finally, we can compute the context specificity factor in this simple model:

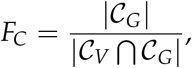

which has a particularly nice interpretation. The inverse, 1/*F*_*C*_, is the proportion of trait-relevant contexts that are affected by the variant. That is, we do not care how many contexts the variant affects, we only care how many contexts are affected *where the gene is relevant to traits*. In this sense *F*_*C*_ captures an intuitive measure of context-specificity where we measure context-specificity weighted by how important each context is to any trait.

This simple model makes a number of strong assumptions. We can relax the assumption that all of the effects of the variant on the contexts are either *β* or 0 and the assumption that the effects of the gene in each context on the corresponding trait is either *γ* or 0. In particular, we will now consider a model where there is still a one-to-one context-to-trait mapping, but the effect of the variant on context *c* is *β*_*c*_ and the effect of the gene in context *c* on trait *c* is *γ*_*c*_ (note that we only need a single index on *γ* now because determining the context determines the trait and *vice versa*). We show this model in Figure S22.

**Figure S22:**
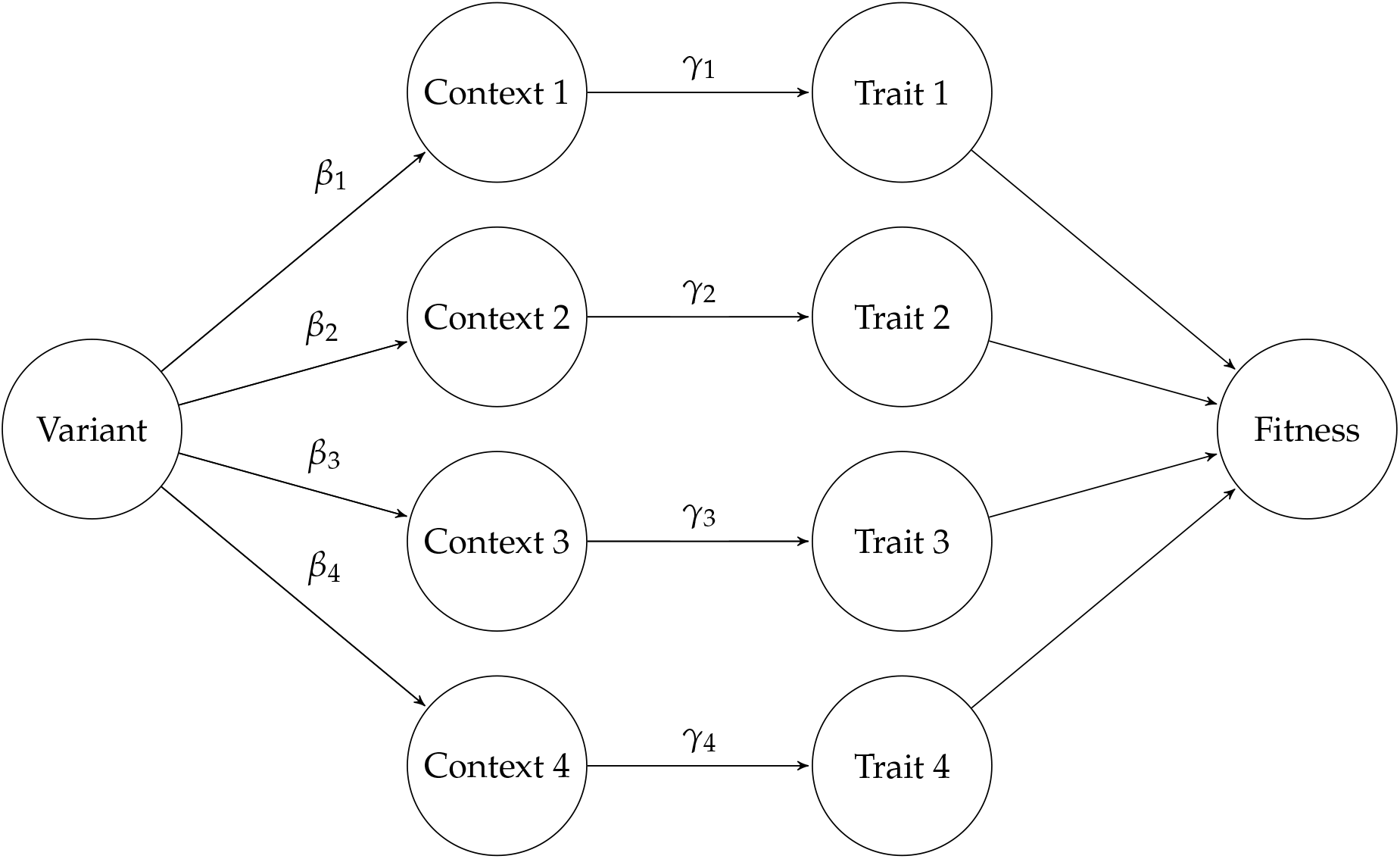
A slight relaxation of the simplest model. There is still a one-to-one correspondence between contexts and traits, but now variants can have arbitrary effects on contexts, and genes can have arbitrary effects on the corresponding trait in each context.

Under this model, we then have

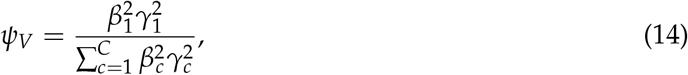

and

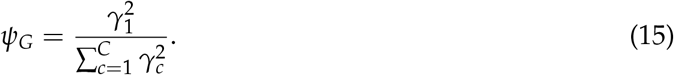

Substituting these into equation 13, we obtain

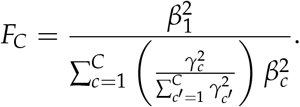

This equation is interpretable — it is the effect of the variant on the focal context relative to a weighted average affect across all contexts where the weights for each context are proportional to how much the gene affects a trait in each context.

In both simplified models, the context specificity factor tracks how much a variant affects the context relevant for the focal trait relative to how much it affects all contexts (weighted by their trait relevance). In either case, these models formalize the intuition that by only affecting a subset of contexts, a variant can be substantially more trait specific than the gene through which it acts.

### B.1 Variants can be less trait specific than the genes through which they act

Our full model (Figure S20) can result in counterintuitive cases where a variant is *less* trait specific than the gene through which it acts. Indeed, even the simplified model shown in Figure S22 can result in cases where *ψ*_*V*_ < *ψ*_*G*_. Since this simplified model is a sub-model of the full model, it implies that such examples are feasible in the full model as well. As a numerical example, assume there are two contexts and take *β* = (1/2, 2) and *γ* = (2, 1/2). Plugging these values into Equations 14 and 15 we obtain

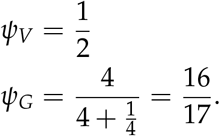

Here we see that the variant evenly affects both traits (and hence is quite non-specific) while the gene very heavily favors the focal trait (and hence is quite specific).

An intuitive explanation for what has happened is that the variant has strong effects on relatively unimportant contexts and weak effects on important contexts. These balance out, and so suddenly contexts that were at very different scales of impact at the gene level become more similar in impact at the variant level. For example, core genes for one trait may have weak effects on other traits when expressed in the wrong context [11]. A variant that hardly effects the important context, but massively perturbs an unimportant context might bring these effects more in line with each other and hence be less specific than the gene.

## C The impact of stabilizing selection on allele frequency in the context of trait specificity and genetic drift

In Appendix A, we derived our results under an assumption of mutation-selection balance, assuming that the strength of selection acting on a variant was proportional to the sum of its squared effects across all traits, 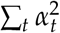. In this Appendix, we extend these results to the case of mutation-selection-drift balance, obtaining the results of Appendix A as a limiting case.

Before beginning, we note that under our model, the effective strength of selection against heterozygotes is 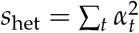, with the fitness in homozygotes being 1. Meanwhile, we defined the trait specificity of the variant to be 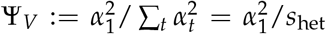, where we assume *α*_1_ is the effect on the focal trait. As such, specifying two of *s*_het_, Ψ_*V*_, or 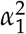 determines the third. In particular,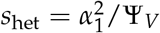.

As discussed in Appendix A, the expected strength of association in GWAS is closely related to the heritability contributed by the variant,

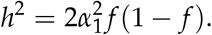

To understand the typical contribution of a variant, we need to compute the expected value of 2 *f* (1 − *f*), which depends on *s*_het_.

It has long been well-understood [38, 61, 63] that under our stabilizing selection setup, the evolutionary dynamics of an allele are approximately equivalent to those of an underdominant model, where individuals homozygous for either allele have fitness 1, but heterozygous individuals have fitness 1 − *s*_het_. We note that under a model of stabilizing selection, heterozygous individuals are not actually less fit than homozygotes on average — it is something of a mathematical coincidence that the evolutionary dynamics under stabilizing selection are equivalent to such a model.

In an underdominant model, the allelic dynamics are described by the stochastic differential equation (SDE)

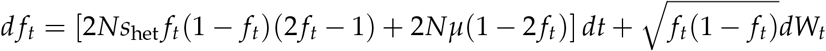

where *N* is the effective population size, *f*_*t*_ is the frequency at time *t, W*_*t*_ is a Wiener process (Brownian motion), and time is measured in units of 2*N* generations. The properties of the stationary distribution of this SDE are well-understood [38,61,63]. In particular, it has been shown that under the stationary distribution of this model,

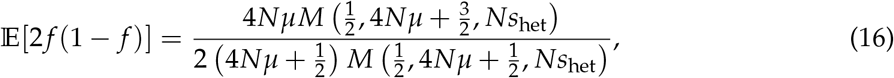

where *M*(., ., .) is the confluent hypergeometric function. In practice, this can be accurately computed numerically for small to moderate values of *Ns*_het_ using `scipy.special.hyp1f1` [84] in Python or `BAS::hypergeometric1F1` [85] in R. For large values of *Ns*_het_ we found these functions to be numerically unstable. For large values of *Ns*_het_, we rely on the asymptotic approximation of the confluent hypergeometric function [86, (13.7.1)],

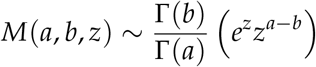

where Γ is Gamma function. After some algebra and noting that Γ(*x* + 1)/Γ(*x*) = *x*, this results in

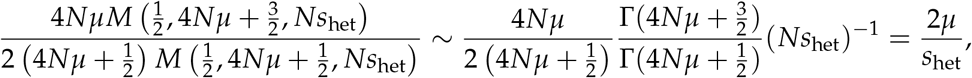

exactly recovering the expectation of 2 *f* (1 − *f*) under mutation-selection balance.

We can also consider the behavior for small values of *s*_het_, where [86, (13.2.2)]

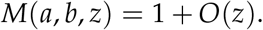

In this case we obtain

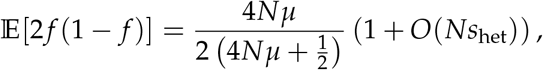

which indicates that the expected value of 2 *f* (1 − *f*) is approximately independent of *Ns*_het_ for values of *Ns*_het_ ≪1. This is consistent with population genetics intuition of these sites being effectively neutral, and hence behaving approximately the same as neutral variants regardless of the precise values of their selection coefficients.

We end with a discussion of the behavior of the expected value of *h*^2^ as a function of 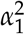 and Ψ_*V*_. To emphasize this reliance of 𝔼*h*^2^ on these quantities, we will write 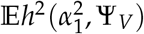. Writing 𝔼*π*(*s*_het_) for 𝔼[2 *f* (1 − *f*)] to emphasize that the expected value of the heterozygosity (*π*) depends on *s*_het_, and recalling that 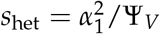, we have

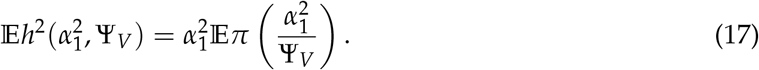

Our above results show that for small 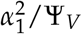, regardless of Ψ_*V*_,

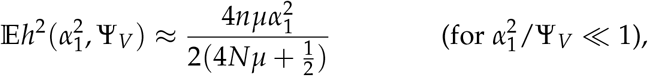

and so expected heritability in this regime is driven essentially solely by the effect size. In contrast, when 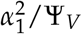 is large, our results imply

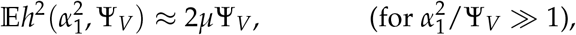

implying that expected strength of association is determined only by Ψ_*V*_. Note that we can be in this regime either by being extremely non-specific 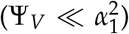 or by having an extremely large effect on the focal trait 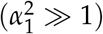.

Finally, using a technical lemma, Lemma C.1, that we prove below, we can show that for any *µ* > 0, *π*(*s*_het_) is a strictly decreasing function of *s*_het_. This result is extremely intuitive as it says that as the effective strength of selection against heterozygotes increases, the expected heterozygosity decreases. This in turn implies that for fixed 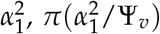 is a strictly *increasing* function of Ψ_*V*_. Finally, if we hold 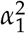 fixed, we see that the only dependence on Ψ_*V*_ in equation 17 is via 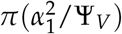. This implies that for fixed 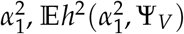 is a strictly increasing function of Ψ_*V*_.

While this result may seem technical, it has a simple interpretation: among all variants with the same trait importance, GWAS will, on average, rank the most trait-specific variants the most highly. Thus, while we focused on the role of trait-specificity in the mutation-selection balance regime in the main text, trait-specificity plays a key role in association study power across the entire range of effect sizes.

### C.1 Technical lemma

#### Lemma C.1.

*Suppose b* ≥ *a* > 0. *Then*,

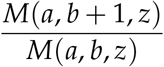

*is a strictly decreasing function of z on z* > 0.

*Proof*. Our proof relies on a technical lemma [87, Lemma 2.1], that states that if all of the following hold:

- *f* (*x*) and *g*(*x*) can be represented as series 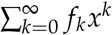 and 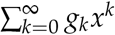 respectively
- Both series converge on −*R* < *x* < *R*
- *g*_*k*_ > 0 for all *k*
- 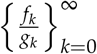 is a decreasing sequence

then *f* (*x*)/*g*(*x*) is a strictly decreasing function of *x* on the interval (0, *R*).

To use this lemma, we use the series representation [86, (13.2.2)],

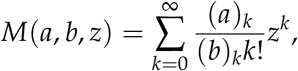

where (*c*)_*k*_ := *c*(*c* + 1) …(*c* + *k* − 1) is the Pochammer symbol, with the convention that (*c*)_0_ := 1. It can be easily shown that this series is convergent for all *z* by noting that (*a*)_*k*_/(*b*)_*k*_ ≤ 1 by assumption that *a* ≤ *b* and comparing terms to the everywhere convergent series for exp *z*. Similar considerations show that *M*(*a, b* + 1, *z*) is everywhere convergent.

We now write

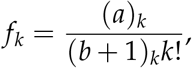

and

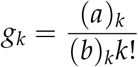

for the coefficients of *z*^*k*^ in the series representations of *M*(*a, b* + 1, *z*) and *M*(*a, b, z*) respectively. To use the lemma, we need to prove that all *g*_*k*_ > 0 and that the ratios *f*_*k*_/*g*_*k*_ are decreasing in *k*. That *g*_*k*_ > 0 for all *k* is trivial, as each *g*_*k*_ is a ratio of positive integers. Now, note that

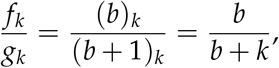

which is strictly decreasing in *k*. Applying [87, Lemma 2.1] completes the proof.

## D Connection between effect sizes and fitness under stabilizing selection

In this Appendix, we discuss the assumptions under which we may assume that the effective strength of selection acting against a variant, *s*_het_, is the sum of that variant’s trait importances,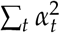.

Our starting point is an arbitrary fitness function, *w*, which maps a vector of trait values, **t**, to a fitness. That is, *w*(**t**) specifies the average fitness of an individual with trait values **t**. We need some mild, technical assumptions on *w* — for our purposes it is sufficient that *w* have an isolated local maximum, and that *w* be twice differentiable at that maximum. Without loss of generality,we may assume that *w*(**t**) has a local maximum at **t** = 0 by redefining **t** as **t** − **t**^∗^ where **t**^∗^ is a local optimum of *w*. We then assume that natural selection is strong enough so that most individuals have traits that are close to the optimum. More precisely, we will assume that the trait values we are interested in are close enough to zero so that terms smaller than 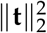 are negligible. As such, we may perform a multivariate Taylor expansion of our fitness function around 0:

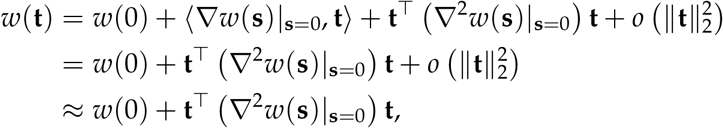

where the second line follows from the fact that 0 is an optimum so the gradient of *w* evaluated at 0 is 0.

Defining **H** := ∇^2^*w*(**s**)|_**s**=0_ as the Hessian (i.e., matrix of second derivatives) of *w* at 0, we obtain that the fitness consequence of having a set of traits **t** is −**t**^⊤^**Ht**. Note that since we are at an isolated local maximum, **H** must be negative definite, which just means that **v**^⊤^**Hv** < 0 for any **v**.

This is exactly the setting of [38] (under certain assumptions discussed below), so we obtain that if a variant causes a change in phenotypes of *α* = (*α*_1_, …, *α*_*T*_) then that results in evolutionary dynamics equivalent to underdominance with a selection coefficient proportional to

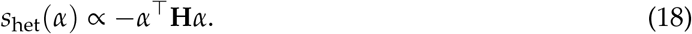

Recently [64], it has been shown that the assumption of independent sites used in [38] to derive equation 18 is incompatible with the Bulmer Effect [62, 88] where variants tend to be in slightly negative LD with other variants that affect the trait in the same direction even if the variants are physically unlinked. Accounting for these subtleties can be well-approximated by changing the constant of proportionality in equation 18, and does not affect our results here [64, equation (43)].

Here we make our first transformation — we absorb the constant of proportionality into *α*. That is, we may scale all of the traits by some constant to make the above proportionality an equality:

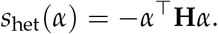

So far we have subtracted a constant from all of our traits and then scaled them by a constant. This does not fundamentally change any of the traits we have measured.

Now, expanding the right-hand side of equation 18 we obtain

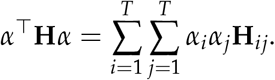

The cross terms make downstream analysis difficult. Furthermore, the constraint that **H** be negative definite implies nontrivial constraints on the values that the entries of **H** may take. To simplify further analysis, we would like to get rid of these cross terms without changing our interpretation of the focal trait (say the first dimension of **t**). The rest of this Appendix will be devoted to accomplishing that task.

Since **H** is negative definite, we may define an inner product by

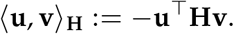

Note that this is different from the usual Euclidean inner product, but we may prove that it is inner product by noting that

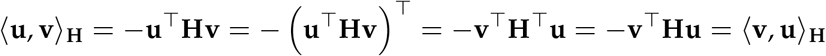

by the symmetry of the Hessian. This shows that our putative inner product is symmetric. Next, note that for scalars *a* and *b*,

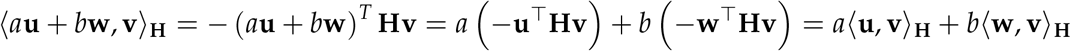

which shows that our putative inner product is linear. Finally for **v**≠0

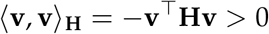

by the negative definiteness of **H** showing that our inner product is positive definite. These three properties define an inner product, so ⟨., .⟩_**H**_ is a bonafide inner product.

Now, since this is inner product, we may perform the Gram-Schmidt process with respect to this inner product starting with the vector (1, 0, 0, …) which corresponds exactly to the focal trait in the original coordinate system. The output of the Gram-Schmidt process is a new basis, *w*_1_, …, *w*_*T*_ such that

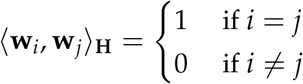

and **w**_1_ ∝ (1, 0, 0, …). These new vectors describe how to change the coordinates of a vector from the original coordinate system to a new coordinate system that is orthonormal with respect to our inner product. In our case, we can think of coordinates as traits. The first coordinate remains unchanged (up to a rescaling that is independent of the other coordinates) from the original coordinate system to the new coordinate system, but the remaining coordinates will be linear combinations of more than one coordinate. This means that in our new coordinate system, we still have the focal trait, but then define new traits in terms of linear combinations of the original traits. By orthonormality, we see that if we write *α* in this new coordinate system,

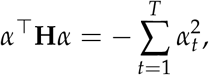

and thus,

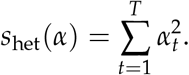

Ultimately, this means that we may define our traits such that the first dimension is a centered and scaled version of the original first trait dimension (centered so its optimum is zero, scaled so that a unit change of the trait *holding all other transformed traits constant* has a unit selection coefficient). The remaining dimensions get scrambled (including with the first original trait dimension), meaning that they are each some linear combination of the original traits.

A subtle point here is that in this coordinate system, the non-focal traits may be in part determined by the focal trait, so we need to be careful in interpreting pleiotropy. For example if we had traits that were weight and 1/height^2^, we could change the coordinate system to instead be weight and BMI. When we talk about a variant that is specific to weight in this new coordinate system, that variant must not affect BMI. But in the original coordinate system, the variant is necessarily not specific to weight because to keep BMI fixed, it must affect both height and weight.

To derive our results, we relied on the argument presented in [38]. That argument requires the trait to be sufficiently polygenic that the genetic component of an individual’s phenotype is approximately Gaussian distributed. It also requires that a new mutation has a random effect, and that that effect is isotropic in our final scaled and rotated trait space. That is, conditioned on its total squared effect, the effect of a new mutation is equally likely to point in any direction of trait space. There are also a number of additional assumptions that are generally met in practice for most traits. Most of these assumptions are either met in practice or do not substantially affect the interpretation of the results. See [38, Supplementary Note Sections 4 and 5].

To summarize, if we choose our coordinate system carefully (i.e., choose he correct set of traits and how to measure them), then under fairly arbitrary models of stabilizing selection we may obtain 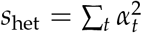. In practice we do not know the fitness function, and so we cannot determine the proper rotation and scaling of trait space to obtain this result. As a result, in the main text, we assume that the traits we consider are such that this result holds. This is obviously a gross approximation, but seems to work surprisingly well in practice, for example in Figure 3**C**. It would be an interesting line of future work to try to estimate **H** (or equivalently learn the proper rotation and scaling of trait space).

## References

[1] Dickinson, M. E., Flenniken, A. M., Ji, X., Teboul, L., Wong, M. D., White, J. K., Meehan, T. F., Weninger, W. J., Westerberg, H., Adissu, H., and others,. High-throughput discovery of novel developmental phenotypes. Nature, 537(7621):508–514, 2016.

[2] Amin, N. D. and Paşca, S. P. Building models of brain disorders with three-dimensional organoids. Neuron, 100(2):389–405, 2018.

[3] Replogle, J. M., Saunders, R. A., Pogson, A. N., Hussmann, J. A., Lenail, A., Guna, A., Masci-broda, L., Wagner, E. J., Adelman, K., Lithwick-Yanai, G., and others,. Mapping information-rich genotype-phenotype landscapes with genome-scale perturb-seq. Cell, 185(14):2559–2575, 2022.

[4] Claussnitzer, M., Cho, J. H., Collins, R., Cox, N. J., Dermitzakis, E. T., Hurles, M. E., Kathiresan, S., Kenny, E. E., Lindgren, C. M., MacArthur, D. G., and others,. A brief history of human disease genetics. Nature, 577(7789):179–189, 2020.

[5] Minikel, E. V., Painter, J. L., Dong, C. C., and Nelson, M. R. Refining the impact of genetic evidence on clinical success. Nature, pages 1–6, 2024.

[6] Finucane, H. K., Bulik-Sullivan, B., Gusev, A., Trynka, G., Reshef, Y., Loh, P.-R., Anttila, V., Xu, H., Zang, C., Farh, K., and others,. Partitioning heritability by functional annotation using genome-wide association summary statistics. Nature genetics, 47(11):1228–1235, 2015.

[7] Calderon, D., Bhaskar, A., Knowles, D. A., Golan, D., Raj, T., Fu, A. Q., and Pritchard, J. K. Inferring relevant cell types for complex traits by using single-cell gene expression. The American Journal of Human Genetics, 101(5):686–699, 2017.

[8] Richard, D., Muthuirulan, P., Young, M., Yengo, L., Vedantam, S., Marouli, E., Bartell, E., Hirschhorn, J., and Capellini, T. D. Functional genomics of human skeletal development and the patterning of height heritability. Cell, 2024.

[9] Schnitzler, G. R., Kang, H., Fang, S., Angom, R. S., Lee-Kim, V. S., Ma, X. R., Zhou, R., Zeng, T., Guo, K., Taylor, M. S., and others,. Convergence of coronary artery disease genes onto endothelial cell programs. Nature, 626(8000):799–807, 2024.

[10] Maurano, M. T., Humbert, R., Rynes, E., Thurman, R. E., Haugen, E., Wang, H., Reynolds, A. P., Sandstrom, R., Qu, H., Brody, J., and others,. Systematic localization of common disease-associated variation in regulatory dna. Science, 337(6099):1190–1195, 2012.

[11] Boyle, E. A., Li, Y. I., and Pritchard, J. K. An expanded view of complex traits: from polygenic to omnigenic. Cell, 169(7):1177–1186, 2017.

[12] Sinnott-Armstrong, N., Naqvi, S., Rivas, M., and Pritchard, J. K. GWAS of three molecular traits highlights core genes and pathways alongside a highly polygenic background. Elife, 10:e58615, 2021.

[13] Yengo, L., Vedantam, S., Marouli, E., Sidorenko, J., Bartell, E., Sakaue, S., Graff, M., Eliasen, A. U., Jiang, Y., Raghavan, S., and others,. A saturated map of common genetic variants associated with human height. Nature, 610(7933):704–712, 2022.

[14] Szustakowski, J. D., Balasubramanian, S., Kvikstad, E., Khalid, S., Bronson, P. G., Sasson, A., Wong, E., Liu, D., Wade Davis, J., Haefliger, C., and others,. Advancing human genetics research and drug discovery through exome sequencing of the UK Biobank. Nature genetics, 53(7):942–948, 2021.

[15] Morgenthaler, S. and Thilly, W. G. A strategy to discover genes that carry multi-allelic or mono-allelic risk for common diseases: a cohort allelic sums test (CAST). Mutation Research/Fundamental and Molecular Mechanisms of Mutagenesis, 615(1-2):28–56, 2007.

[16] Backman, J. D., Li, A. H., Marcketta, A., Sun, D., Mbatchou, J., Kessler, M. D., Benner, C., Liu, D., Locke, A. E., Balasubramanian, S., and others,. Exome sequencing and analysis of 454,787 UK Biobank participants. Nature, 599(7886):628–634, 2021.

[17] Singh, T., Poterba, T., Curtis, D., Akil, H., Al Eissa, M., Barchas, J. D., Bass, N., Bigdeli, T. B., Breen, G., Bromet, E. J., and others,. Rare coding variants in ten genes confer substantial risk for schizophrenia. Nature, 604(7906):509–516, 2022.

[18] Zhou, D., Zhou, Y., Xu, Y., Meng, R., and Gamazon, E. R. A phenome-wide scan reveals convergence of common and rare variant associations. Genome Medicine, 15(1):101, 2023.

[19] Weiner, D. J., Nadig, A., Jagadeesh, K. A., Dey, K. K., Neale, B. M., Robinson, E. B., Karczewski, K. J., and O’Connor, L. J. Polygenic architecture of rare coding variation across 394,783 exomes. Nature, 614(7948):492–499, 2023.

[20] Bycroft, C., Freeman, C., Petkova, D., Band, G., Elliott, L. T., Sharp, K., Motyer, A., Vukcevic, D., Delaneau, O., O’Connell, J., and others,. The UK Biobank resource with deep phenotyping and genomic data. Nature, 562(7726):203–209, 2018.

[21] Tsuji, T. and Kunieda, T. A loss-of-function mutation in natriuretic peptide receptor 2 (Npr2) gene is responsible for disproportionate dwarfism in cn/cn mouse. Journal of Biological Chemistry, 280(14):14288–14292, 2005.

[22] Olney, R. C., Bükülmez, H., Bartels, C. F., Prickett, T. C., Espiner, E. A., Potter, L. R., and Warman, M. L. Heterozygous mutations in natriuretic peptide receptor-B (NPR2) are associated with short stature. The Journal of Clinical Endocrinology & Metabolism, 91(4):1229–1232, 2006.

[23] Vasques, G. A., Amano, N., Docko, A. J., Funari, M. F., Quedas, E. P., Nishi, M. Y., Arnhold, I. J., Hasegawa, T., and Jorge, A. A. Heterozygous mutations in natriuretic peptide receptor-B (NPR2) gene as a cause of short stature in patients initially classified as idiopathic short stature. The Journal of Clinical Endocrinology & Metabolism, 98(10):E1636–E1644, 2013.

[24] Amano, N., Mukai, T., Ito, Y., Narumi, S., Tanaka, T., Yokoya, S., Ogata, T., and Hasegawa, T. Identification and functional characterization of two novel NPR2 mutations in Japanese patients with short stature. The Journal of Clinical Endocrinology & Metabolism, 99(4):E713– E718, 2014.

[25] Wang, S. R., Jacobsen, C. M., Carmichael, H., Edmund, A. B., Robinson, J. W., Olney, R. C., Miller, T. C., Moon, J. E., Mericq, V., Potter, L. R., and others,. Heterozygous mutations in natriuretic peptide receptor-B (NPR2) gene as a cause of short stature. Human mutation, 36(4):474–481, 2015.

[26] Amberger, J. S., Bocchini, C. A., Schiettecatte, F., Scott, A. F., and Hamosh, A. OMIM. org: Online Mendelian Inheritance in Man (OMIM®), an online catalog of human genes and genetic disorders. Nucleic acids research, 43(D1):D789–D798, 2015.

[27] Conery, M. and Grant, S. F. Human height: a model common complex trait. Annals of human biology, 50(1):258–266, 2023.

[28] To, K., Fei, L., Pett, J. P., Roberts, K., Blain, R., Polański, K., Li, T., Yayon, N., He, P., Xu, C., and others,. A multi-omic atlas of human embryonic skeletal development. Nature, 635(8039):657– 667, 2024.

[29] Chuang, P.-T. and McMahon, A. P. Vertebrate Hedgehog signalling modulated by induction of a Hedgehog-binding protein. Nature, 397(6720):617–621, 1999.

[30] Bishop, B., Aricescu, A. R., Harlos, K., O’callaghan, C. A., Jones, E. Y., and Siebold, C. Structural insights into hedgehog ligand sequestration by the human hedgehog-interacting protein HHIP. Nature structural & molecular biology, 16(7):698–703, 2009.

[31] Briscoe, J. and Thérond, P. P. The mechanisms of hedgehog signalling and its roles in development and disease. Nature reviews Molecular cell biology, 14(7):416–429, 2013.

[32] Gillespie, J. H. Population genetics: a concise guide. JHU press, 2004.

[33] Zeng, T., Spence, J. P., Mostafavi, H., and Pritchard, J. K. Bayesian estimation of gene constraint from an evolutionary model with gene features. Nature Genetics, 56(8):1632–1643, 2024.

[34] Sanjak, J. S., Sidorenko, J., Robinson, M. R., Thornton, K. R., and Visscher, P. M. Evidence of directional and stabilizing selection in contemporary humans. Proceedings of the National Academy of Sciences, 115(1):151–156, 2018.

[35] Sella, G. and Barton, N. H. Thinking about the evolution of complex traits in the era of genome-wide association studies. Annual Review of Genomics and Human Genetics, 20(Volume 20, 2019):461–493, 2019.

[36] Patel, R. A., Weiss, C. L., Zhu, H., Mostafavi, H., Simons, Y. B., Spence, J. P., and Pritchard, J. K. Conditional frequency spectra as a tool for studying selection on complex traits in biobanks. bioRxiv, pages 2024–06, 2024.

[37] Koch, E., Connally, N., Baya, N., Reeve, M., Daly, M., Neale, B., Lander, E., Bloemendal, A., and Sunyaev, S. Genetic association data are broadly consistent with stabilizing selection shaping human common diseases and traits. bioRxiv, pages 2024–06, 2024.

[38] Simons, Y. B., Bullaughey, K., Hudson, R. R., and Sella, G. A population genetic interpretation of GWAS findings for human quantitative traits. PLoS biology, 16(3):e2002985, 2018.

[39] O’Connor, L. J., Schoech, A. P., Hormozdiari, F., Gazal, S., Patterson, N., and Price, A. L. Extreme polygenicity of complex traits is explained by negative selection. The American Journal of Human Genetics, 105(3):456–476, 2019.

[40] Zou, Z., Ohta, T., and Oki, S. ChIP-Atlas 3.0: a data-mining suite to explore chromosome architecture together with large-scale regulome data. Nucleic Acids Research, page gkae358, 2024.

[41] Uhlén, M., Fagerberg, L., Hallström, B. M., Lindskog, C., Oksvold, P., Mardinoglu, A., Sivertsson, Å., Kampf, C., Sjöstedt, E., Asplund, A., and others,. Tissue-based map of the human proteome. Science, 347(6220):1260419, 2015.

[42] Gazal, S., Finucane, H. K., Furlotte, N. A., Loh, P.-R., Palamara, P. F., Liu, X., Schoech, A., Bulik-Sullivan, B., Neale, B. M., Gusev, A., and others,. Linkage disequilibrium–dependent architecture of human complex traits shows action of negative selection. Nature genetics, 49(10):1421–1427, 2017.

[43] Karczewski, K. J., Francioli, L. C., Tiao, G., Cummings, B. B., Alföldi, J., Wang, Q., Collins, R. L., Laricchia, K. M., Ganna, A., Birnbaum, D. P., and others,. The mutational constraint spectrum quantified from variation in 141,456 humans. Nature, 581(7809):434–443, 2020.

[44] Pickrell, J. K., Berisa, T., Liu, J. Z., Ségurel, L., Tung, J. Y., and Hinds, D. A. Detection and interpretation of shared genetic influences on 42 human traits. Nature genetics, 48(7):709–717, 2016.

[45] Watanabe, K., Stringer, S., Frei, O., Umicévić Mirkov, M., de Leeuw, C., Polderman, T. J., van der Sluis, S., Andreassen, O. A., Neale, B. M., and Posthuma, D. A global overview of pleiotropy and genetic architecture in complex traits. Nature genetics, 51(9):1339–1348, 2019.

[46] Qi, G., Chhetri, S. B., Ray, D., Dutta, D., Battle, A., Bhattacharjee, S., and Chatterjee, N. Genome-wide large-scale multi-trait analysis characterizes global patterns of pleiotropy and unique trait-specific variants. Nature Communications, 15(1):6985, 2024.

[47] Mostafavi, H., Spence, J. P., Naqvi, S., and Pritchard, J. K. Systematic differences in discovery of genetic effects on gene expression and complex traits. Nature Genetics, 55(11):1866–1875, 2023.

[48] Milind, N., Smith, C. J., Zhu, H., Spence, J. P., and Pritchard, J. K. Buffering and non-monotonic behavior of gene dosage response curves for human complex traits. medRxiv, pages 2024–11, 2024.

[49] Weiner, D. J., Gazal, S., Robinson, E. B., and O’Connor, L. J. Partitioning gene-mediated disease heritability without eQTLs. The American Journal of Human Genetics, 109(3):405–416, 2022.

[50] Pathan, N., Deng, W. Q., Di Scipio, M., Khan, M., Mao, S., Morton, R. W., Lali, R., Pigeyre, M., Chong, M. R., and Paré, G. A method to estimate the contribution of rare coding variants to complex trait heritability. Nature Communications, 15(1):1245, 2024.

[51] Lappalainen, T., Li, Y. I., Ramachandran, S., and Gusev, A. Genetic and molecular architecture of complex traits. Cell, 187(5):1059–1075, 2024.

[52] Akbari, P., Gilani, A., Sosina, O., Kosmicki, J. A., Khrimian, L., Fang, Y.-Y., Persaud, T., Garcia, V., Sun, D., Li, A., and others,. Sequencing of 640,000 exomes identifies GPR75 variants associated with protection from obesity. Science, 373(6550):eabf8683, 2021.

[53] Gazal, S., Weissbrod, O., Hormozdiari, F., Dey, K. K., Nasser, J., Jagadeesh, K. A., Weiner, D. J., Shi, H., Fulco, C. P., O’Connor, L. J., and others,. Combining snp-to-gene linking strategies to identify disease genes and assess disease omnigenicity. Nature Genetics, 54(6):827–836, 2022.

[54] Freund, M. K., Burch, K. S., Shi, H., Mancuso, N., Kichaev, G., Garske, K. M., Pan, D. Z., Miao, Z., Mohlke, K. L., Laakso, M., and others,. Phenotype-specific enrichment of mendelian disorder genes near GWAS regions across 62 complex traits. The American Journal of Human Genetics, 103(4):535–552, 2018.

[55] Umans, B. D., Battle, A., and Gilad, Y. Where are the disease-associated eQTLs? Trends in Genetics, 37(2):109–124, 2021.

[56] Burch, K. S., Hou, K., Ding, Y., Wang, Y., Gazal, S., Shi, H., and Pasaniuc, B. Partitioning gene-level contributions to complex-trait heritability by allele frequency identifies disease-relevant genes. The American Journal of Human Genetics, 109(4):692–709, 2022.

[57] de Leeuw, C. A., Mooij, J. M., Heskes, T., and Posthuma, D. MAGMA: generalized gene-set analysis of GWAS data. PLoS computational biology, 11(4):e1004219, 2015.

[58] Duffy, Á., Petrazzini, B. O., Stein, D., Park, J. K., Forrest, I. S., Gibson, K., Vy, H. M., Chen, R., Márquez-Luna, C., Mort, M., and others,. Development of a human genetics-guided priority score for 19,365 genes and 399 drug indications. Nature Genetics, 56(1):51–59, 2024.

[59] Mbatchou, J., Barnard, L., Backman, J., Marcketta, A., Kosmicki, J. A., Ziyatdinov, A., Benner, C., O’Dushlaine, C., Barber, M., Boutkov, B., and others,. Computationally efficient whole-genome regression for quantitative and binary traits. Nature genetics, 53(7):1097–1103, 2021.

[60] Berisa, T. and Pickrell, J. K. Approximately independent linkage disequilibrium blocks in human populations. Bioinformatics, 32(2):283, 2016.

[61] Bulmer, M. G. The genetic variability of polygenic characters under optimizing selection, mutation and drift. Genetics Research, 19(1):17–25, 1972.

[62] Bulmer, M. G. Linkage disequilibrium and genetic variability. Genetics Research, 23(3):281– 289, 1974.

[63] Keightley, P. D. and Hill, W. G. Quantitative genetic variability maintained by mutation-stabilizing selection balance in finite populations. Genetics Research, 52(1):33–43, 1988.

[64] Negm, S. and Veller, C. The effect of long-range linkage disequilibrium on allele-frequency dynamics under stabilizing selection. bioRxiv, pages 2024–06, 2024.

[65] Zhu, X. and Stephens, M. Bayesian large-scale multiple regression with summary statistics from genome-wide association studies. The annals of applied statistics, 11(3):1561, 2017.

[66] Zeng, T., Spence, J. P., Mostafavi, H., and Pritchard, J. K. s_het estimates from genebayes and other supplementary datasets, December 2023.

[67] Quinlan, A. R. and Hall, I. M. BEDTools: a flexible suite of utilities for comparing genomic features. Bioinformatics, 26(6):841–842, 2010.

[68] Barrett, T., Wilhite, S. E., Ledoux, P., Evangelista, C., Kim, I. F., Tomashevsky, M., Marshall, K. A., Phillippy, K. H., Sherman, P. M., Holko, M., and others,. NCBI GEO: archive for functional genomics data sets—update. Nucleic acids research, 41(D1):D991–D995, 2012.

[69] Hicks, M. R., Hiserodt, J., Paras, K., Fujiwara, W., Eskin, A., Jan, M., Xi, H., Young, C. S., Evseenko, D., Nelson, S. F., and others,. ERBB3 and NGFR mark a distinct skeletal muscle progenitor cell in human development and hPSCs. Nature cell biology, 20(1):46–57, 2018.

[70] Ferguson, G. B., Van Handel, B., Bay, M., Fiziev, P., Org, T., Lee, S., Shkhyan, R., Banks, N. W., Scheinberg, M., Wu, L., and others,. Mapping molecular landmarks of human skeletal ontogeny and pluripotent stem cell-derived articular chondrocytes. Nature communications, 9(1):3634, 2018.

[71] Consortium, I. H. . and others,. Integrating common and rare genetic variation in diverse human populations. Nature, 467(7311):52, 2010.

[72] McLaren, W., Gil, L., Hunt, S. E., Riat, H. S., Ritchie, G. R., Thormann, A., Flicek, P., and Cunningham, F. The ensembl variant effect predictor. Genome biology, 17:1–14, 2016.

[73] Morales, J., Pujar, S., Loveland, J. E., Astashyn, A., Bennett, R., Berry, A., Cox, E., Davidson, C., Ermolaeva, O., Farrell, C. M., and others,. A joint NCBI and EMBL-EBI transcript set for clinical genomics and research. Nature, 604(7905):310–315, 2022.

[74] Spence, J. P., Zeng, T., Mostafavi, H., and Pritchard, J. K. Scaling the discrete-time Wright– Fisher model to biobank-scale datasets. Genetics, 225(3):iyad168, 2023.

[75] Simons, Y. B., Mostafavi, H., Smith, C. J., Pritchard, J. K., and Sella, G. Simple scaling laws control the genetic architectures of human complex traits. bioRxiv, pages 2022–10, 2022.

[76] Bulik-Sullivan, B. K., Loh, P.-R., Finucane, H. K., Ripke, S., Yang, J., of the Psychiatric Genomics Consortium, S. W. G., Patterson, N., Daly, M. J., Price, A. L., and Neale, B. M. LD Score regression distinguishes confounding from polygenicity in genome-wide association studies. Nature genetics, 47(3):291–295, 2015.

[77] Gamazon, E. R., Wheeler, H. E., Shah, K. P., Mozaffari, S. V., Aquino-Michaels, K., Carroll, R. J., Eyler, A. E., Denny, J. C., Consortium, G., Nicolae, D. L., and others,. A gene-based association method for mapping traits using reference transcriptome data. Nature genetics, 47(9):1091–1098, 2015.

[78] Gusev, A., Ko, A., Shi, H., Bhatia, G., Chung, W., Penninx, B. W., Jansen, R., De Geus, E. J., Boomsma, D. I., Wright, F. A., and others,. Integrative approaches for large-scale transcriptome-wide association studies. Nature genetics, 48(3):245–252, 2016.

[79] Privé, F., Arbel, J., and Vilhjálmsson, B. J. LDpred2: better, faster, stronger. Bioinformatics, 36(22-23):5424–5431, 2020.

[80] Wang, G., Sarkar, A., Carbonetto, P., and Stephens, M. A simple new approach to variable selection in regression, with application to genetic fine mapping. Journal of the Royal Statistical Society Series B: Statistical Methodology, 82(5):1273–1300, 2020.

[81] Spence, J. P., Sinnott-Armstrong, N., Assimes, T. L., and Pritchard, J. K. A flexible modeling and inference framework for estimating variant effect sizes from GWAS summary statistics. BioRxiv, pages 2022–04, 2022.

[82] Good, B. H. Linkage disequilibrium between rare mutations. Genetics, 220(4):iyac004, 2022.

[83] Dominici, D. E. The inverse of the cumulative standard normal probability function. Integral Transforms and Special Functions, 14(4):281–292, 2003.

[84] Virtanen, P., Gommers, R., Oliphant, T. E., Haberland, M., Reddy, T., Cournapeau, D., Burovski, E., Peterson, P., Weckesser, W., Bright, J., and others,. Scipy 1.0: fundamental algorithms for scientific computing in python. Nature methods, 17(3):261–272, 2020.

[85] Clyde, M. BAS: Bayesian Variable Selection and Model Averaging using Bayesian Adaptive Sampling, 2024. R package version 1.7.5.

[86] NIST Digital Library of Mathematical Functions. https://dlmf.nist.gov/, Release 1.2.2 of 2024-09-15. F. W. J. Olver, A. B. Olde Daalhuis, D. W. Lozier, B. I. Schneider, R. F. Boisvert, C. W. Clark, B. R. Miller, B. V. Saunders, H. S. Cohl, and M. A. McClain, eds.

[87] Ponnusamy, S. and Vuorinen, M. Asymptotic expansions and inequalities for hypergeometric function. Mathematika, 44(2):278–301, 1997.

[88] Bulmer, M. G. The effect of selection on genetic variability. The American Naturalist, 105(943):201–211, 1971.

